# Restoring Antibiotic Sensitivity via Metabolic Pathway Manipulation in *Pseudomonas aeruginosa*

**DOI:** 10.64898/2026.10.01.755869

**Authors:** Hannah M. Doherty, Jennifer van der Laan, Daniel Martínez-Martínez, Theresa Fink, Katharina Stanzel, Huda Ahmad, Leonie Helfer, Ivan Andrew, Laurence Game, Iliyana Kaneva, Alex Montoya, Holger Kramer, João Mokochinski, Susanne Brodesser, Ignacio Sottorff, Erwin Bohn, Tobias Bollenbach, Manuel Banzhaf, Filipe Cabreiro

## Abstract

Stopping antimicrobial multidrug resistance (AMR) is essential to maintain the efficacy of current treatments and sustain modern medicine, but current control relies on a coordinated One Health approach involving stewardship, infection prevention and other control measures, without direct actionable mechanisms against resistance itself. Here, integrating proteomics and metabolomics with high-throughput *Caenorhabditis elegans* infection models, we show that unique synergy of biotin biosynthesis inhibition with metformin uncouples virulence from bacterial viability shutting down major *Pseudomonas aeruginosa* (PA) virulence mechanisms that impair host survival. Under directed evolution induced by central carbon metabolic stress, laboratory and clinical PA strain isolates acquire mutations in the MexAB-OprM multidrug efflux pump, leading to reversal of the AMR phenotype and restored susceptibility to multiple legacy and last-resort antibiotics such as meropenem. Our findings reveal a previously unexploited vulnerability in PA, its co-dependence on biotin biosynthesis and energy metabolism for virulence and resistance. This work establishes a conceptual framework in which metabolic co-targeting can abolish pathogenesis and drive reversal of antimicrobial resistance. It lays the foundation for first-in-class, non-antibiotic PA-selective anti-virulence and resistance-reversing therapies, key to extending the lifesaving duration of existing antibiotics.

## MAIN

Antimicrobial efficacy is an integral pillar within modern medicine, being essential for successful eradication and prevention of infection worldwide. In 2021, a global study associated 4.71 million deaths and directly attributed 1.14 million deaths to AMR ^1^. The pathogen *Pseudomonas aeruginosa* (PA) was ranked globally as the 4^th^ deadliest AMR organism when carbapenem resistant, and the 2^nd^ deadliest within high-income countries ^2^. Given its propensity to cause both acute and chronic infections within immunocompromised people, PA is highly problematic in the nosocomial environment. The arsenal of virulence factors, and intrinsic and acquired resistance mechanisms this pathogen encodes, makes it a formidable pathogen ^3,4^. PA is renowned for forming difficult-to-eradicate biofilms on hospital equipment, as well producing toxins and host-degrading enzymes which cause extensive cellular damage ^3,5,6^. Furthermore, rapidly rising antimicrobial resistance (AMR) is leaving once treatable infections with increased mortality rates ^1^. Therefore, research and development of new treatment options is paramount. As such, shifting focus to repurposing currently approved therapeutics, or to the development of drugs with new mechanisms of action could hold significant potential. Historically, the development of antibiotics has been primarily focused on inhibiting bacterial growth via the targeting of DNA, RNA, protein or cell wall biosynthesis. Unfortunately, AMR has been evolved against all antibiotics targeting these functions, making the discovery of new targets pivotal. A considerably underexplored drug target, which holds significant potential, is central metabolism ^7,8^. For example, the regulation of central metabolism must be carefully controlled by bacteria in response to their environment, balancing energy consumption and production to ensure survival ^8^. One drug, prime for repurposing, is the anti-glycaemic type-2 diabetes drug metformin, due to its discovered impact on bacterial metabolism and its proven safety with over 50 years of abundant use ^9–12^. Several studies have explored metformin’s suitability for repurposing across a variety of contexts, from anti-cancer, anti-inflammatory, anti-aging, and anti-tuberculosis, all with promising evidence ^11,13,14^. Further to this, anti-virulence studies have shown that metformin may supress virulence factor production ^15,16^, however the mechanisms involved have not been exploited.

Here, we bridge this gap in knowledge by combining high-throughput drug-microbe-host infection phenotyping with omics approaches. By exposing *C. elegans* to metformin and growing on individual clones of transposon functional libraries of PA, we identify biotin as an essential micronutrient mediating virulence through metabolic rewiring of central carbon and energy metabolism under metformin. Further, we discovered that synergy between biotin biosynthesis inhibition and metformin treatment drives evolution towards dysfunctional MexAB-OprM efflux pumps. Remarkably, loss of MexAB-OprM function leads to AMR reversal due to collateral sensitivity (CS) to common, as well as last resource antibiotics in PA clinical isolates and slowed down newly acquired resistance evolution within *P. aeruginosa* UCBPP-PA14 (PA14).

### *C. elegans* survival with PA by metformin is mediated by biotin

To test whether metformin can sufficiently suppress PA pathogenesis and improve host health for infection clearance, we utilised *Caenorhabditis elegans* as an infection model. The detrimental nature of PA to *C. elegans* survival is known to be both strain and nutrient dependent ^17^. Furthermore, similar to mouse infections, PA killing of worms is largely facilitated by the production of virulence factors, specifically siderophores ^17,18^. As metformin-mediated beneficial host effects are largely impacted by nutrient composition ^9^ we evaluated the survival of *C. elegans* wildtype N2 in the presence of PA14 and metformin on different nutritional contexts. Survival was tested on three base media with varying levels of nutritional strength: standard nematode growth media (NGM), slow killing (SK) medium, and LB-NGM (Extended Data Fig. 1a). A significant increase in survival was observed in the presence of metformin on NGM media (p < 0.001) and SK medium (p < 0.001) but not in LB-NGM (Fig. 1a, Extended Data Table 1), suggesting a nutrient dependent response to metformin modulation of *C. elegans* survival on PA. To understand how metformin and nutrient-mediated changes to host mechanisms affected *C. elegans* survival, we performed bulk RNA sequencing. Metformin caused a change in several enriched Gene Ontology (GO) molecular functions (including the innate immune response and defence responses to other organisms) in NGM media but not LB-NGM, further supporting the survival phenotypes (Fig. 1b, Extended Data Table 2). To identify the mechanisms by which nutrition regulates PA14 pathogenesis in *C. elegans* we leveraged our transcription data to design a high-throughput screen of a PA14 transposon-insertion library in the presence of metformin in LB-NGM (Fig. 1c) ^19^. For the selection of an appropriate biosensor capturing bacterial-drug-nutrient changes, we selected worm genes upregulated in the presence of metformin in a nutrient dependent manner and narrowed the selection to *gpdh-1*, *sod-3*, *fzo-1*, *argk-1* and *irg-6* (Extended Data Fig. 1b-f, Extended Data Table 3)*. Argk-1,* an arginine kinase functionally described as a downstream effector through S6 kinase of the mTOR pathway ^20^, previously shown to be up-regulated in response to metformin in the presence of *E*. *coli* ^9^, now showed a significant upregulation in the presence of metformin in a nutrient dependent manner with PA14 (Extended Data Fig. 1f, Extended Data Table 3). The high-throughput screen revealed multiple gene disruptions which increased the expression of *argk-1* with metformin (Fig. 1d, Extended Data Table 2). Of note, the PA14 Tn::*gacA* was revealed, which is a global response regulator associated with virulence in *C. elegans* and mammals (Fig. 1d, Extended Data Fig. 1g, Extended Data Table 2) ^17,21^. To further understand the pathways linked to this phenotype, we performed an enrichment analysis on the top 2% of genes with the highest upregulation of *argk-1.* A total of 38 enriched pathways were found, and those with a fold enrichment higher than 5 were highlighted (Fig. 1e, Extended Data Table 2). Among the pathways with the highest fold enrichment were the glycerol-3-phosphate dehydrogenase complex, biotin biosynthesis, pyrimidine biosynthesis and RNA binding domain, implying an overall role of these PA14 pathways in metformin effects in a nutrient-dependent manner. To investigate if these transcriptional changes predicted survival phenotypes, we measured *C. elegans* N2 survival in the presence of *P. aeruginosa* mutants with and without metformin in LB-NGM media, nutrient conditions where no survival differences were observed for metformin treatment. Overall, we found that the inhibition of several PA14 pathways, such as pyrimidine and sulphur metabolism, increased worm survival even in the absence of metformin with some exceptions (Extended Data Fig. 2a-n, Extended Data Table 1). Metformin marginally but significantly improved the survival of *C. elegans* N2 with PA14 Tn::*glpD* by 5% (±7%, Cox proportional hazards, CPH, p < 0.05). More strikingly, disruption in biotin biosynthesis, PA14 Tn::*bioA*, showed a large significant synergy with metformin to extend *C. elegans* survival by 99% (±19%, CPH p < 0.001, Extended Data Fig. 2o-p, Extended Data Table 1). Out of all the candidate pathways, this indicated that biotin bioavailability may explain the nutrient-dependent signature of metformin in PA14 survival.

The significance of these findings was further confirmed via a metformin dose-response survival assay. Metformin significantly improved *C. elegans* survival in the presence of PA14 Tn::*bioF* in a dose-dependent manner (CPH p < 0.001, Fig. 1f, Extended Data Table 1). Furthermore, this synergistic effect on survival between biotin depletion and metformin was consistent for all genes within the biotin biosynthesis pathway (CPH p < 0.001, Extended Data Fig. 3a-d, Extended Data Table 1). Supplementation of 400 nM of biotin abolished the rescued survival phenotype of worms exposed to PA14 Tn::*bioF* and metformin, confirming that the loss of biotin production was responsible for the metformin-associated increase in survival (Fig. 1g, Extended Data Table 1). Lastly, unlike worms in PA14 or biotin mutants, which show typical signs of tissue decay upon infection, the morphology of *C. elegans* with different biotin biosynthesis mutants plus metformin showed improved worm size despite extensive bacterial gut colonisation, visually showing improvement of *C. elegans* cell integrity and overall health (Fig. 1h, Extended Data Fig. 3e, Extended Data Table 4). To understand how PA14 biotin biosynthesis inhibition synergises with metformin to increase *C. elegans* survival, we performed bulk RNA sequencing. Worms exposed to Tn::*gacA* were included as an avirulent control condition. As predicted, the transcriptome of worms exposed to PA14 Tn::*bioF* and metformin shifted towards similar expression patterns as worms on Tn::*gacA*. Worms exposed to Tn::*bioF* or metformin independently showed transcriptomes similar to PA14 WT, supporting a synergy between biotin biosynthesis depletion and metformin in host survival (Fig. 1i). In addition, relative transcript levels of key genes including *argk-1* showed similar trends as observed for metformin in NGM media (Extended Data Fig. 3f, Extended Data Table 4). Metformin-specific upregulation of *argk-1* with biotin depletion was further confirmed through fluorescence imaging of transgenic P*argk-1*::GFP worms grown on individual mutants of the biotin biosynthesis pathway (Extended Data Fig. 3g-h, Extended Data Table 4). Altogether, this suggests that the increased survival of worms may be due to decreased virulence from the biotin-deficient mutants in the presence of metformin.

**Figure 1:**
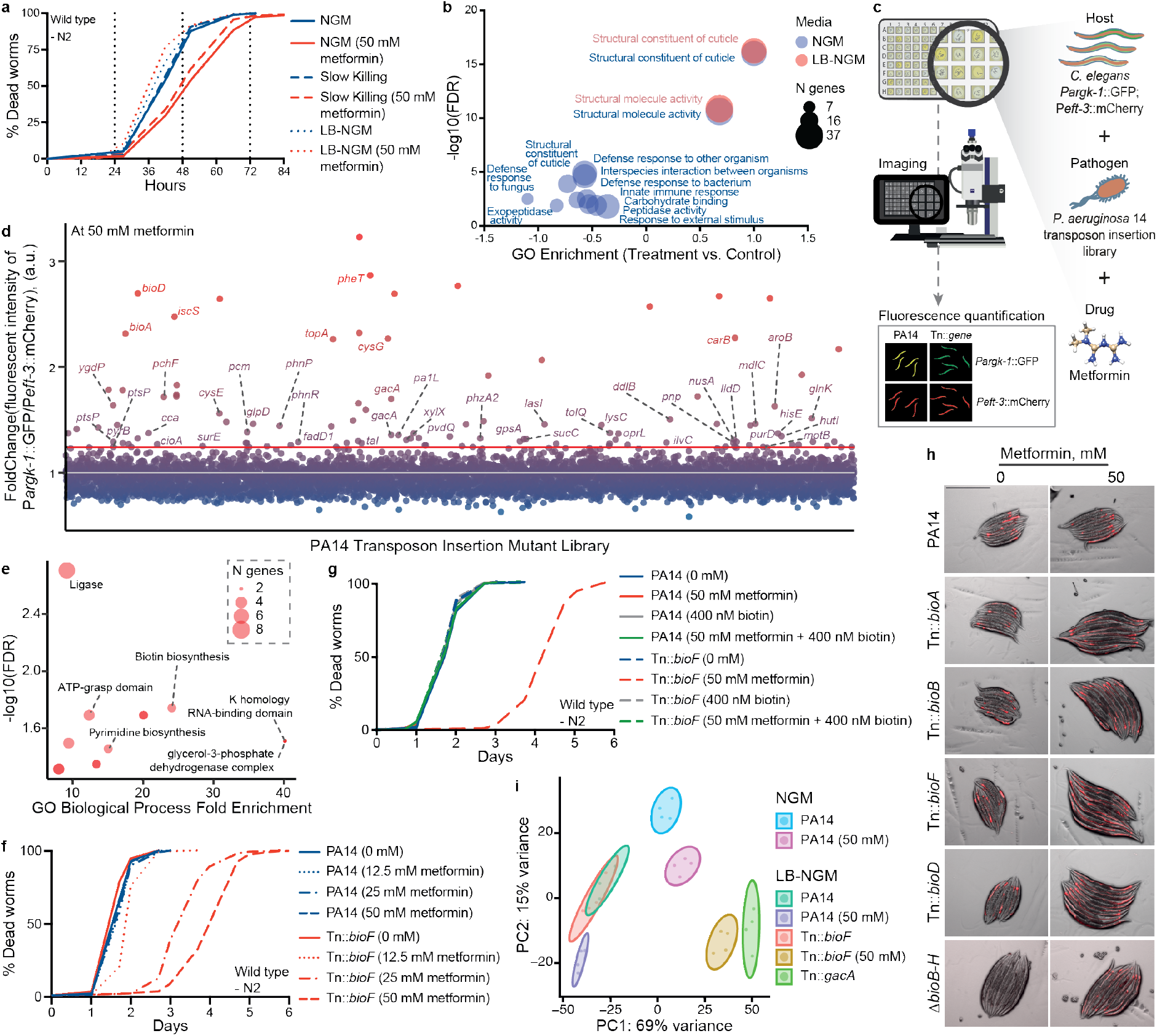
Metformin synergises with biotin depletion to improve *C. elegans* survival with PA14. **a,** Influence of media nutrition level on survival of *C. elegans* N2 infected with PA14, without (blue) and with 50 mM metformin (red). Metformin extension of survival is nutrient dependent (*n* = 279-394). **b,** Transcriptome Gene Ontology enrichment, categories Molecular Function and Biological Processes, of up and downregulated *C. elegans* genes on metformin (50 mM) vs. no metformin LB-NGM or NGM media and infected with PA14 (*n* = 4). Total enrichment data is provided in Extended Data Table 2. **c,** Methodology overview used to screen PA14 transposon insertion mutants which induce increased P*argk-1*::GFP fluorescence with metformin (50 mM). **d,** Manhattan plot of PA14 transposon insertion mutants which resulted in increased P*argk-1*::GFP fluorescence compared to WT PA14 within *C. elegans*. Highlighted (above red line) are 2% of PA14 transposon insertion mutants activating P*argk-1*::GFP fluorescence the highest with metformin. Unannotated dots are uncharacterised genes to date. **e,** Gene Ontology Biological Process enrichment of top 2% genes which when disrupted in PA14 increased P*argk-1*::GFP fluorescence. Shown are enriched categories with a fold change > 5. Total enrichment data is provided in Extended Data Table 2. **f,** PA14 Tn::*bioF* showed a metformin-concentration dependent increase in survival of *C. elegans* N2 (*n* = 242-378). **g,** Environmental biotin addition abrogates metformin-dependant survival increase for *C. elegans* infected with PA14 Tn::*bioF* (*n* = 277-449). All survival statistics are provided in Extended Data Table 1. **h,** Visualisation of pMF440 in viable PA14 and respective mutants in *C. elegans* intestine, with and without metformin exposure. Worms showed morphological improvements in the presence of PA14 biotin depleted mutants and metformin. Scale bar reflects 1000 µm. **i,** Principal component analysis of the *C. elegans* N2 transcriptomes after exposure to PA14 compared to biotin deficient mutants with and without metformin on LB-NGM (*n* = 4).

**Extended Data Figure 1:**
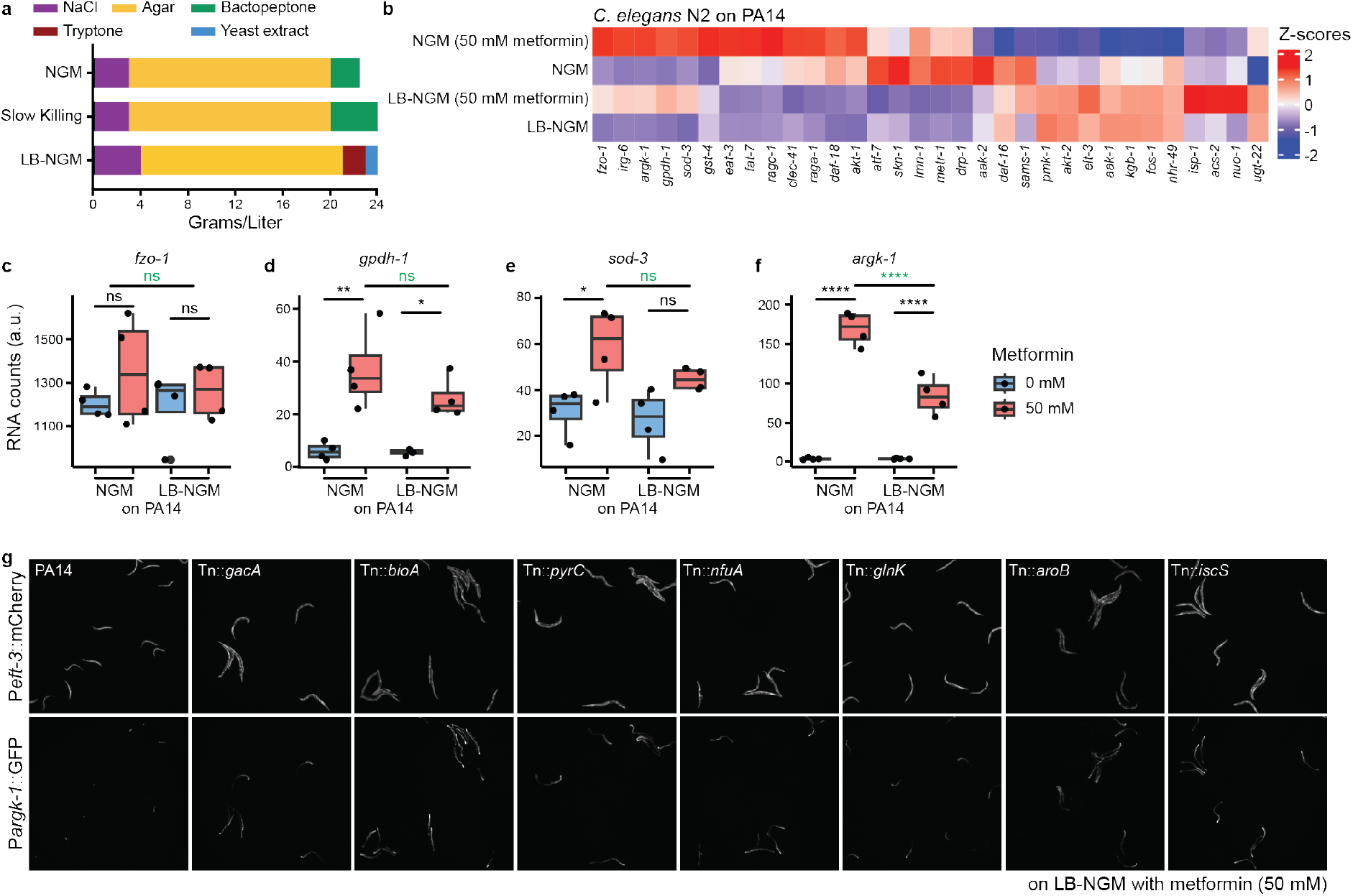
Metformin and nutrient deprivation were able to improve *C. elegans* survival with PA14. **a,** Nutrient composition of different media, prior to cholesterol, MgSO_4_, CaCl_2_ and KH_2_PO_4_ addition, to determine *C. elegans* survival with metformin influenced by nutrient availability. **b,** Z-scores of relative RNA counts per gene for a selection of *C. elegans* N2 genes involved in stress response and regulation, innate immunity, longevity, mitochondrial homeostasis and respiration, without and with metformin (50 mM) on NGM or LB-NGM (*n* = 4). **c-f**, RNA counts of *fzo-1, gpdh-1, sod-3* and *argk-1* in *C. elegans* N2, exposed to PA14 on NGM or LB-NGM, were upregulated in the presence of metformin (*n* = 4). All statistics are provided in Extended Data Table 3. **g**, Example images of *C. elegans* P*argk-1*::GFP;P*eft-3*::mCherry on PA14 wildtype and selected PA14 transposon insertion mutant hits on LB-NGM supplemented with metformin (50 mM).

**Extended Data Figure 2:**
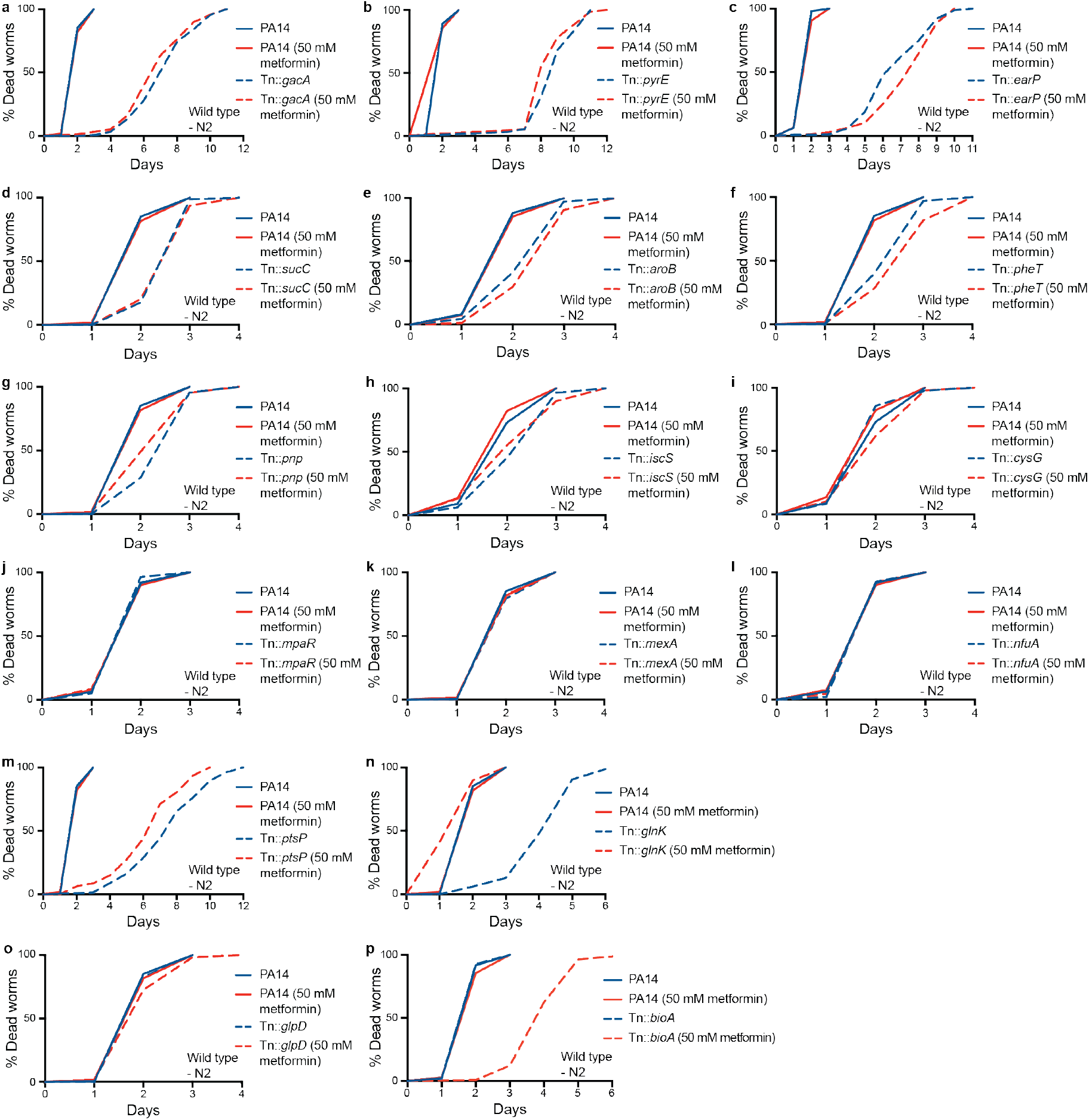
Disruption in metabolic pathways of PA14 altered *C. elegans* N2 survival in metformin dependent or independent manners. **a-i,** Survival of *C. elegans* N2 with and without metformin (50 mM) with disruption: in the global response regulator PA14 Tn::*gacA* (p < 0.0001, *n* = 151-154) (**a**), the orotate phosphoribosyltransferase Tn::*pyrE* (p < 0.0001, *n* = 78-91) (**b**), the glycosyltransferase Tn::*earP* (p < 0.0001, *n* = 207-306) (**c**), the succinyl synthetase Tn::*sucC* (p < 0.0001, *n* = 86-116) (**d**), the 3-hydroquinate synthase Tn::*aroB* (p < 0.0001, *n* = 151-241) (**e**), the phenylalanyl-tRNA synthetase Tn::*pheT* (p < 0.0001, *n* = 73-84) (**f**), the polyribonucleotide nucleotidyl transferase Tn::*pnp* (p < 0.0001, *n* = 68-69) (**g**), the L-cysteine desulferase Tn::*iscS* (p < 0.0001, *n* = 69-113) (**h**), and the siroheme synthase Tn::*cysG* (p < 0.01, *n* = 89-123) (**i**). **j-l**, Disruption of the PA14 PQS regulator Tn::*mpaR* (*n* = 151-251) (**j**), Tn::*mexA,* part of the multi-drug MexAB-OprM efflux pump (*n* = 194-208) (**k**), and Tn::*nfuA,* encoding for iron-sulfur cluster delivery protein (*n* = 126-130)(**l**). **m-n**, Mutation of the pyocyanin production regulator, Tn::*ptsP* (p < 0.01, *n* = 130-136) (**m**) or the nitrogen regulator, PA14 Tn::*glnK* (p < 0.001, *n* = 148-195) (**n**). **o**, Survival in glycerol-3-dehydrogenase PA14 Tn::*glpD* mutants (p < 0.05, *n* = 178-183). **p**, Metformin significantly synergised with disruption in biotin biosynthesis, PA14 Tn::*bioA* (p < 0.001, *n* = 88-160). All p values represent CPH results. All survival statistics are provided in Extended Data Table 1.

**Extended Data Figure 3:**
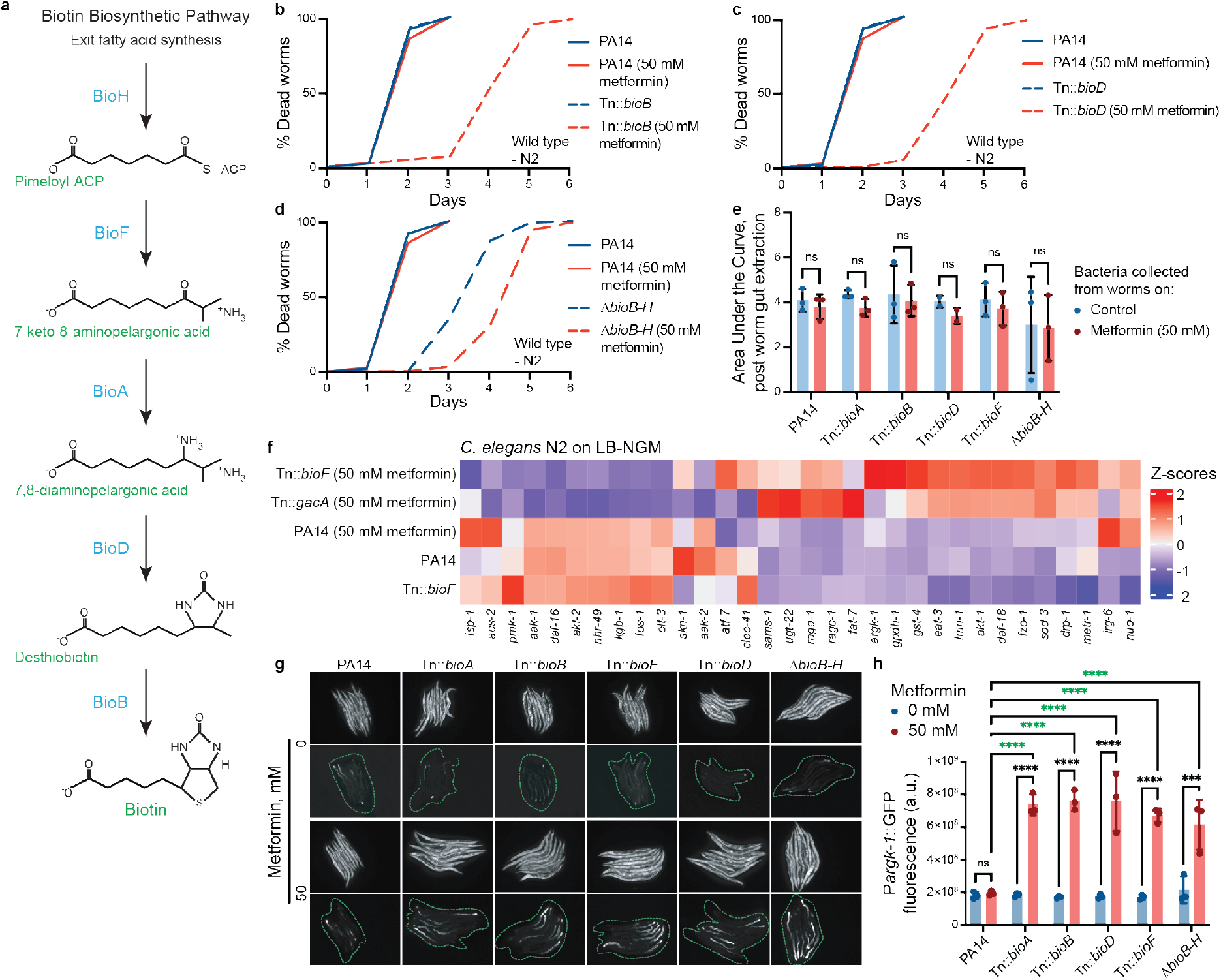
Biotin deficient PA14 and metformin showed a reduction in pathogenesis in *C. elegans*. **a,** Biotin biosynthesis pathway within PA. **b-c,** PA14 Tn::*bioB* (CPH p < 0.0001, *n* = 91-147) and Tn::*bioD* (p < 0.0001, *n* = 113-229) were able to significantly improve *C. elegans* N2 survival with 50 mM metformin. **d,** PA14 Δ*bioB-H* improved *C. elegans* survival with and without addition of metformin (CPH p < 0.0001, *n* = 107-148). All survival statistics are provided in Extended Data Table 1. **e,** Area under the curve levels representing bacterial growth showed viable bacteria extracted from *C. elegans* intestines on LB-NGM with and without metformin (ns: p > 0.05, *n* = 2-3). Statistics are provided in Extended Data Table 4. **f,** Z-scores of relative RNA counts per gene for a selection of *C. elegans* N2 genes, including the arginine kinase *argk-1* (*n* = 4). **g-h,** P*argk-1*::GFP fluorescent levels were significantly upregulated with all PA14 biotin biosynthesis depletion mutants and metformin (****: p < 0.0001). (*n* = 4). Statistics are provided in Extended Data Table 4.

### Metformin affects central metabolic pathways inhibiting growth and virulence in biotin-deficient PA mutants

Since no reduction in bacterial load was observed within infected worms, despite significant increased survival, a potential loss in virulence was tested. Firstly, we assessed bacterial fitness within increasing concentrations of metformin. A dose-dependent response was observed for both wildtype PA14 and the biotin-pathway mutants (Fig. 2a, Extended Data Fig. 4a, Extended Data Table 5 and 6). Critically, this difference was abolished with the supplementation of exogenous biotin, confirming biotin’s role in metformin effects. Secondly, we performed proteomics for PA14 and PA14 Tn::*bioD* with exposure to metformin. An enrichment analysis of the KEGG categories for significantly up- and down-regulated proteins revealed dramatic changes in the proteome of PA14 Tn::*bioD* but not in the WT PA14, with striking differences in the down-regulation of proteins associated with virulence associated pathways, namely phenazine biosynthesis, biofilm formation, and quorum sensing (Fig. 2b, Extended Data Fig. 4d,e, Extended Data Table 5 and 6). Conversely, proteins involved in the biosynthesis of siderophores, were upregulated within metformin treated PA14 Tn::*bioD*. Further inspection of the proteins in this category revealed that the pyochelin biosynthesis proteins (PchA/B/C/D/F/G) specifically, were upregulated (Extended Data Fig. 4f, Extended Data Table 6). Interestingly, although not enriched due to the absence of the pathway in KEGG, proteins involved the biosynthesis and uptake of the primary siderophore, pyoverdine, were downregulated with metformin. These included PvdD/F/N/O/P/Q and FpvA/B, with log2(Fold Change) values between -1.06 and -5.53 (Extended Data Fig. 4g, Extended Data Table 6), indicating a shift away from the production of the primary siderophore with metformin and biotin biosynthesis inhibition. To confirm that the changes in the levels of virulence-associated proteins resulted in changes to virulence-associated metabolite production, we performed untargeted metabolomics ^22^. Despite the upregulation of pyochelin synthesis proteins, the metabolomic analysis revealed a reduction in pyochelin abundance within metformin treated PA14 Tn::*bioF* (Fig. 2f, Extended Data Table 5). This may be the result of the FptA Fe(III)-pyochelin receptor being down regulated (Log2(FC): -1.92) for Tn::*bioD* treated with metformin (Extended Data Fig. 4f, Extended Data Table 6). Overall, this suggests that re-uptake of iron-bound pyochelin may be reduced, despite a potential increase in production. Concurrent with the proteomics analysis was the observed reduction in phenazine metabolites, namely 1-phenazinol and pyocyanin (Fig. 2d,e, Extended Data Table 5). The untargeted metabolomics further revealed a reduction in quorum sensing, with multiple quinolones being reduced in PA14 Tn::*bioF* in the presence of metformin. The overall reduction in virulent metabolites of PA14 Tn::*bioF* with metformin revealed similar improvements to the avirulent PA14 Tn::*gacA*, confirming the mechanism behind the reduction in pathogenesis (Fig. 2c, Extended Data Table 5). Complementary to the P*argk-1*::GFP screen and survival assays, these data together suggest that the combination of biotin biosynthesis inhibition plus metformin results in reduced virulence and pathogenicity in PA despite similar infection levels.

**Figure 2:**
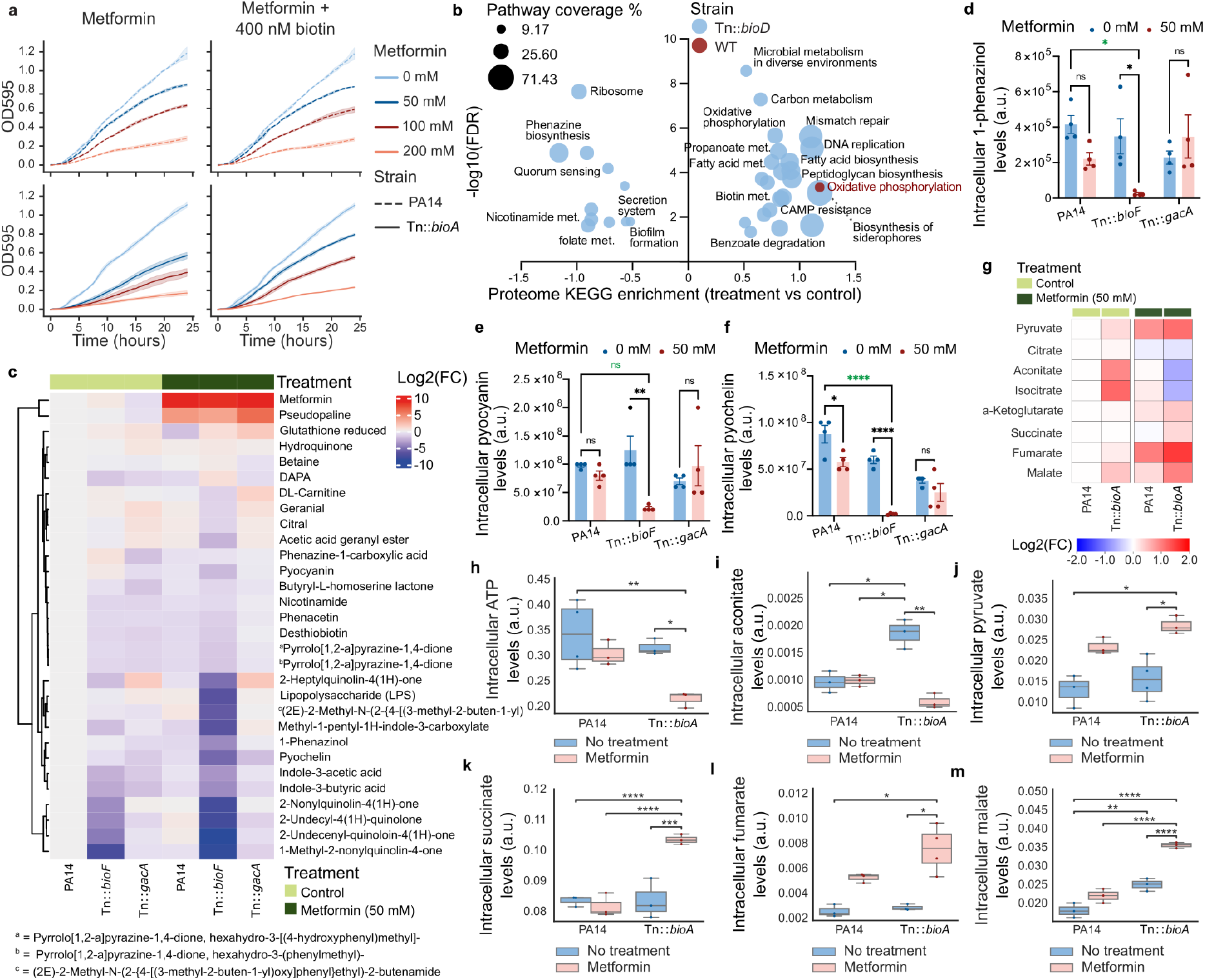
Metformin affects PA14 growth, energy metabolism, and virulence factor production with biotin synthesis loss. **a**, Growth curves of PA14 and PA14 biotin synthesis mutants in increasing concentrations of metformin, with and without biotin supplementation (*n* = 3). **b,** KEGG enrichment of up- and down-regulated proteins within WT PA14 (red) and PA14 Tn::*bioD* (blue) in the presence of metformin (*n* = 5). **c,** Log2(FC) expression of metabolites within PA14, PA14 Tn::*gacA*, and PA14 Tn::*bioF*, with and without metformin in comparison to PA14 no metformin control. (*n* = 4). **d-f,** Intracellular levels of 1-phenazinol (**d**), pyocyanin (**e**), and pyochelin (**f**) within PA14, PA14 Tn::*gacA*, and PA14 Tn::*bioF*, with (grey) and without (red) 50 mM metformin. (ns: p > 0.05, *: 0.01 < p <= 0.05, **: 0.001< p <= 0.01, ***: 0.0001 < p <= 0.001, ****: p <= 0.0001). (*n* = 4). **g,** Log2(FC) of TCA cycle intermediates for PA14, and PA14 Tn::*bioA* with and without metformin vs PA14 control. **h-m**, Intracellular levels of ATP (**h**), aconitate (**i**), pyruvate (**j**), succinate (**k**), fumarate (**l**), and malate (**m**). (ns: p > 0.05, *: 0.01 < p <= 0.05, **: 0.001< p <= 0.01, ***: 0.0001 < p <= 0.001, ****: p <= 0.0001). (*n* = 3-4, indicated by dot points). All statistics are provided in Extended Data Table 5.

**Extended Data Figure 4:**
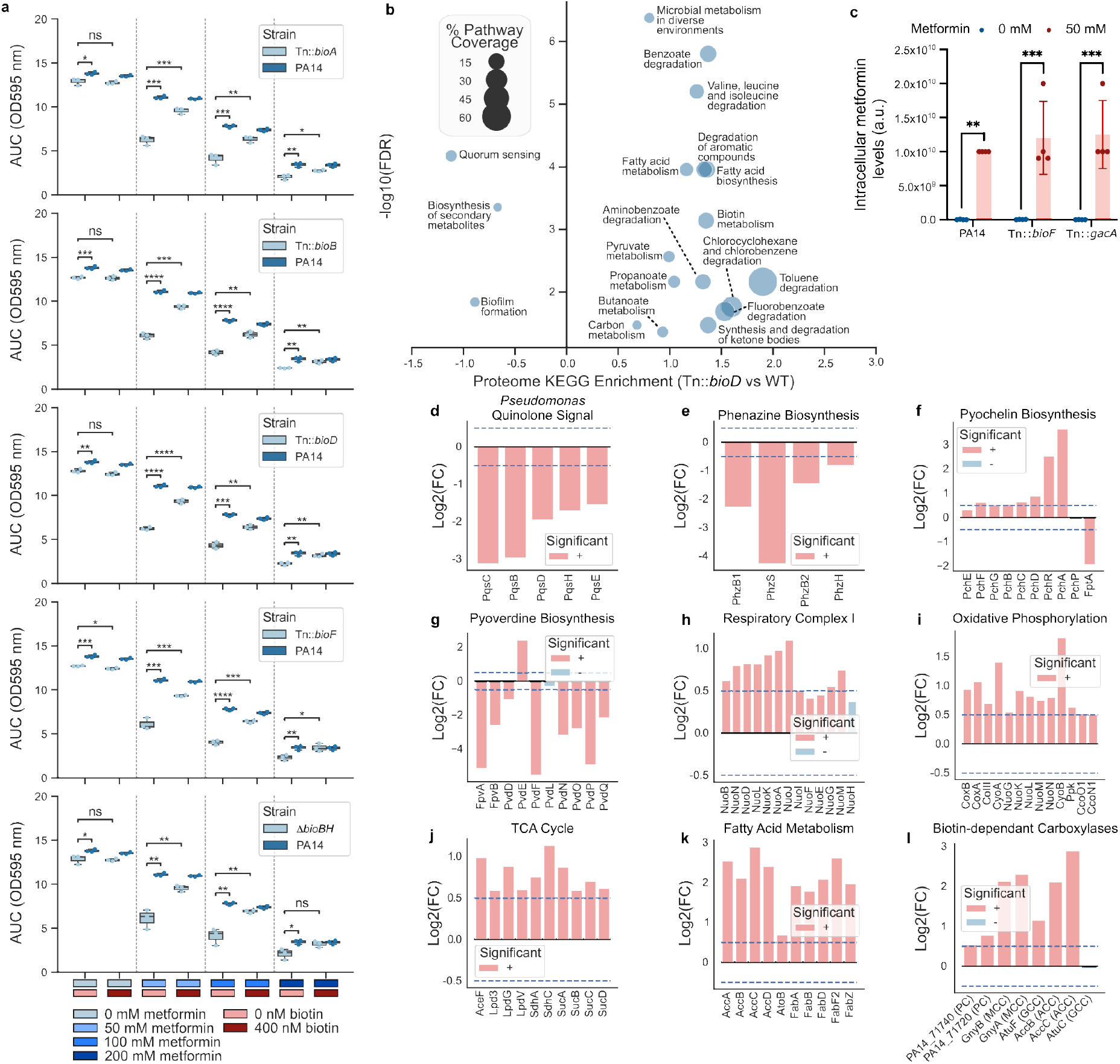
Metformin affects PA14 growth, energy metabolism, and virulence factor production with biotin synthesis loss. **a,** Area under the curve (AUC) values for growth curves of PA14 and PA14 biotin synthesis mutants in increasing concentrations of metformin and supplementation with biotin. (ns: p > 0.05, *: 0.01 < p <= 0.05, **: 0.001< p <= 0.01, ***: 0.0001 < p <= 0.001, ****: p <= 0.0001). (*n* = 3). **b,** KEGG enrichment of up- and down-regulated proteins within WT PA14 and PA14 Tn::*bioD*. (*n* = 5). **c,** Intracellular levels of metformin within PA14, PA14 Tn::*gacA*, and PA14 Tn::*bioF*, with (grey) and without (red) 50 mM metformin treatment. (ns: p > 0.05, *: 0.01 < p <= 0.05, **: 0.001< p <= 0.01, ***: 0.0001 < p <= 0.001, ****: p <= 0.0001). (*n* = 4). **d-k,** Log2(FC) values, for PA14 Tn::*bioD* 50 mM metformin vs PA14 Tn::*bioD* control, of *pseudomonas* quinolone signal proteins **d**), phenazine biosynthesis proteins (**e**), pyochelin synthesis associated proteins (**f**), pyoverdine synthesis related proteins (**g**), Respiratory complex I (NDH-I) proteins (**h**), oxidative phosphorylation proteins (**i**), TCA cycle related enriched proteins (**j**), fatty acid metabolism enriched proteins (**k**) and biotin-dependant carboxylases (**l**) from the proteomic analysis. Blue dashed lines represent the thresholds for inclusion within the enrichment analysis. (*n* = 5). All statistics are provided in Extended Data Table 6.

Biotin plays a key metabolic role within PA via its function as a co-factor for multiple biotin-dependent carboxylases. This is clearly recapitulated by the enrichment of proteins from biotin-associated pathways when comparing proteins upregulated between PA14 and PA14 Tn::*bioD* in the absence of metformin (Extended Data Fig. 4b,l, Extended Data Table 6). Many of these pathways, including fatty acid metabolism/biosynthesis; benzoate degradation; carbon metabolism; valine, leucine and isoleucine degradation; and pyruvate metabolism were further upregulated and enriched with metformin, supporting the synergistic interaction between biotin depletion and metformin treatment (Fig. 2b, Extended Data Table 5).

One pathway upregulated in both PA14 and PA14 Tn::*bioD* treatment groups was oxidative phosphorylation (Fig. 2b, Extended Data Table 5). Significantly, within this group were multiple constituents of the NDH-1 NADH dehydrogenase, NuoB/G/I-N for Tn::*bioD* and NuoI/N/H within PA14 (Extended Data Fig. 4h, Extended Data Table 6). Notably, NDH-1 is homologous to mitochondrial respiratory complex I within the electron transport chain (ETC), a proposed inhibition-target of metformin in humans ^13^. Also upregulated within the oxidative phosphorylation pathway for both WT and Tn::*bioD* with metformin treatment were CoIII, the cytochrome oxidase C subunit III, and subunits of succinate dehydrogenase, SdhD (WT) and SdhA/C (Tn::*bioD*). Despite their similar enrichment, increased pathway coverage was observed for Tn::*bioD* (16/57 for Tn::*bioD* vs. 6/57 for WT) (Extended Data Fig. 4i, Extended Data Table 6). The ETC is integral for the generation of the proton motive force (PMF) and thus ATP generation. Consequently, the overexpression of multiple complexes within the ETC within metformin treated Tn::*bioD* suggests an attempt to re-establish the PMF or increase ATP generation.

Since ETC-associated protein overexpression is heightened with biotin biosynthesis inhibition, biotin restriction must be involved in the increased need for ETC function. To confirm this, we performed targeted metabolomics for anionic metabolites. Critically, intracellular ATP levels were significantly reduced within PA14 Tn::*bioA* with metformin exposure but not wildtype PA14 (Fig. 2h, Extended Data Table 5), revealing that energetic disruption is occurring with biotin synthesis inhibition and metformin. Furthermore, multiple TCA cycle intermediates equally showed altered abundance in this condition, consistent with the TCA cycle enrichment observed in the proteomics data (Extended Data Fig. 4j, Extended Data Table 6). Here, pyruvate, succinate, fumarate and malate accumulated, whereas, citrate, cis-aconitate and isocitrate showed decreased levels (Fig. 2g,i-m, Extended Data Table 5) suggesting a block within the TCA cycle leading to metabolic intermediates accumulation and limitation. Loss of biotin synthesis and metformin treatment could therefore lead to a two-front disruption of oxidative phosphorylation, via decreases in electron donors from the TCA cycle and inhibition of NDH-1, potentially resulting in an dramatic disruption in energy generation, as indicated by the reduced ATP levels and changes in TCA metabolites.

### Metformin resistance yields multidrug collateral sensitivity

Since metformin treatment of PA14 limited in biotin production imposes a drastic metabolic pressure and rewiring, we tested if these conditions would lead to the development of resistance. Incubation of biotin-deficient mutants for several days in the presence of metformin gave rise to suppressor mutant colonies for which we observed higher *in vitro* metformin resistance (Extended Data Fig. 5a, Extended Data Table 8). Metformin resistance in PA14 Tn::*bioA* and Tn::*bioF* abolished prolonged survival of *C. elegans* N2 in the presence of metformin, likely due to higher levels of virulent metabolites such as phenazines (Extended Data Fig. 5b-d, Extended Data Table 1 and 8). To understand where in the genome the resistance-conferring mutations had arisen, we sequenced metformin-resistant (MR) colonies and their parents (Fig. 3a, Extended Data Table 7). For all MR mutants, mutations were identified in components of the major efflux pump MexAB-OprM. Since many AMR bacteria, including PA, enhance antibiotic resistance through efflux pump overexpression^23^, the functional consequences of these mutations were examined using AlphaFold 3 structural predictions ^24^. Three different mutations were present in MexA: (i), a mis-sense mutation (Val129Leu; PA14 Tn::*bioA* MR1.3), analogous to previously reported Val to Met substitutions known to disrupt MexA-OprM interactions ^25^; (ii), an in-frame deletion mutation of Leu193-Pro195 (PA14 Tn::*bioA* MR2.2) (Fig. 3b); and (iii) a premature stop codon at Glu357 truncating residues 358-383 (PA14 Tn::*bioF* MR1.2). Two mutations were identified in MexB, including a mis-sense substitution in Arg971Trp (PA14 Tn::*bioA* MR1.1) and an in-frame deletion spanning Leu724-Ser750 (PA14 Tn::*bioA* MR1.2, PA14 Tn::*bioA* MR2.1) (Fig. 3b). A single frameshift mutation in OprM (from Ser169) rendered the downstream structure unresolvable by AlphaFold, suggesting complete loss of function (Fig. 3b). Collectively, these alterations are predicted to disrupt the structural integrity of the MexAB-OprM complex, resulting in efflux pump inactivation as a direct consequence from the selective pressure imposed by metformin and biotin synthesis inhibition on cellular energetics. Despite this, ATP levels did not increase between metformin exposed PA14 Tn::*bioA* MR1.3 and PA14 Tn::*bioA*, suggesting that ATP may be redistributed with loss of efflux pump function, allowing survival (Extended Data Fig. 5e, Extended Data Table 8). However, alterations within TCA cycle intermediates were observed (Extended Data Fig. 5f, Extended Data Table 8). Reversions to WT levels were observed for malate and succinate, for metformin-treated PA14 Tn::*bioA* MR1.3 (Extended Data Fig. 5g,h, Extended Data Table 8). Reduction in these intermediates, suggests an alleviation of TCA cycle blockage with MexAB-OprM disruption.

**Figure 3:**
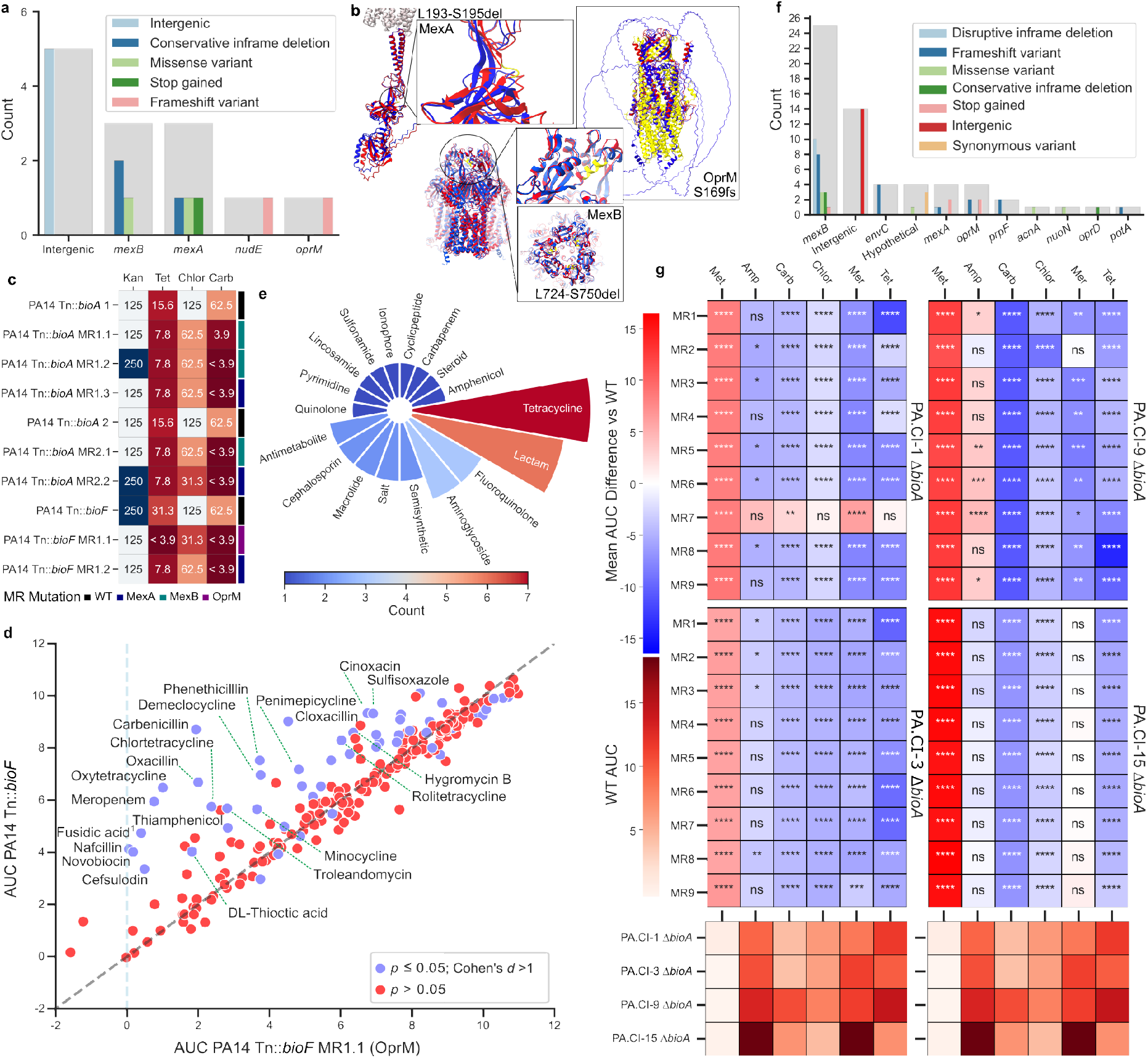
Mutations within the MexAB-OprM efflux pump induce metformin resistance with concurrent multidrug collateral sensitivity. **a,f,** Count of suppressor mutations for different genes and the type of mutation for PA14 (**a**) and clinical isolate (**f**) mutants. Total counts are represented by the grey bar. **b**, AlphaFold structures of WT (red and yellow) MexA (left), MexB (center), and OprM (right) overlayed with the mutated structures (blue). Yellow portions represent the regions within the WT where the mutations occur. **c**, MIC for PA14 Tn::*bioA*, PA14 Tn::*bioF* and their metformin resistant (MR) mutants. (*n* = 3). **d**, BIOLOG chemical sensitivity screen comparing the average area under the curve (AUC) for the growth of PA14 Tn::*bioF* and PA14 Tn::*bioF* MR1.1 for all replicates within all concentrations of different chemical perturbations. Labelled points have an absolute AUC difference > 2. ^1^ = Fusidic acid sodium salt. (*n* = 3). **e,** Counts of antibiotic types within the enrichment category ‘Antibiotics’ for the BIOLOG screen. **g,** Difference in AUC between MR mutants and parents to metformin and MexAB-OprM antibiotic substrates (blue-red scale). Parent AUC in same conditions (white-red scale). Amp: Ampicillin, Carb: Carbenicillin, Chlor: Chloramphenicol, Kan: Kanamycin, Mer: Meropenem, Met: Metformin, Tet: Tetracycline. (ns: p > 0.05, *: 0.01 < p <= 0.05, **: 0.001< p <= 0.01, ***: 0.0001 < p <= 0.001, ****: p <= 0.0001). (*n* = 3). All statistics are provided in Extended Data Table 7.

To further assess if these mutations disrupted MexAB-OprM function, we measured the MICs for carbenicillin, tetracycline, chloramphenicol and kanamycin for the MR mutants and their parental strains. For all MR mutant strains, a reduction of at least 2-fold compared to their respective parents was seen for carbenicillin, tetracycline and chloramphenicol, three known substrates of the MexAB-OprM pump ^26^ (Fig. 3c, Extended Data Table 7). An increase in sensitivity to kanamycin, which is not effluxed by MexAB-OprM, was only observed within the PA14 Tn::*bioF* MR mutants. For PA14 Tn::*bioF* MR1.1, this is likely due to OprM involvement within the MexXY-OprM efflux pump, whose substrate specificity includes aminoglycosides ^26^. These data support AlphaFold 3 predictions and the functional impairment of MexAB-OprM.

Given the increased sensitivity to carbenicillin, tetracycline and chloramphenicol, it stood to reason that the MR mutants may be susceptible to other substrates of the pump. Therefore, we utilised Biolog bacterial chemical sensitivity assay plates to test PA14 Tn::*bioF* MR1.1 sensitivity to a broad range of compounds, including multiple pump substrates. The area under the curve (AUC) for growth in each condition was averaged across all concentrations and PA14 Tn::*bioF* MR1.1 was compared to PA14 Tn::*bioF* (Fig. 3d, Extended Data Table 7). A subset of compounds inhibited the growth of PA14 Tn::*bioF* MR1.1 significantly more than the metformin-sensitive parent. This subset contained multiple antibiotic classes, including penicillins, cephalosporins, carbapenems, tetracyclines, macrolides, aminoglycosides, fluoroquinolones, sulphonamides, and novobiocin (Fig. 3d, Extended Data Table 7). Compounds which induced statistically significant differences in growth with an absolute Cohen’s *d* value greater than 1 were subjected to an enrichment analysis for the compound type. The study revealed a significant enrichment for antibiotic compounds, with tetracyclines also showing independent enrichment (Extended Data Fig. 6a, Extended Data Table 9). Although only tetracyclines were enriched, multiple classes of antibiotics were included within the enriched ‘Antibiotic’ category (Fig. 3e, Extended Data Table 7). To further test MR mutant sensitivity, PA14 Tn::*bioA* MR1.1, MR1.2 and MR1.3, were exposed to a subset of Biolog plates. As with PA14 Tn::*bioF* MR1.1, increased sensitivity was found to multiple antibiotics, despite the reduced compound array. For all PA14 Tn::*bioA* MR mutants, increased sensitivity was gained for beta-lactams, sulfathiazole, novobiocin and nalidixic acid (Extended Data Fig. 6b-d, Extended Data Table 9). These data confirm that metformin-induced loss of function of the efflux pump MexAB-OprM leads to increased antibiotic sensitivity against a wide range of antimicrobial compounds.

**Extended Data Figure 5:**
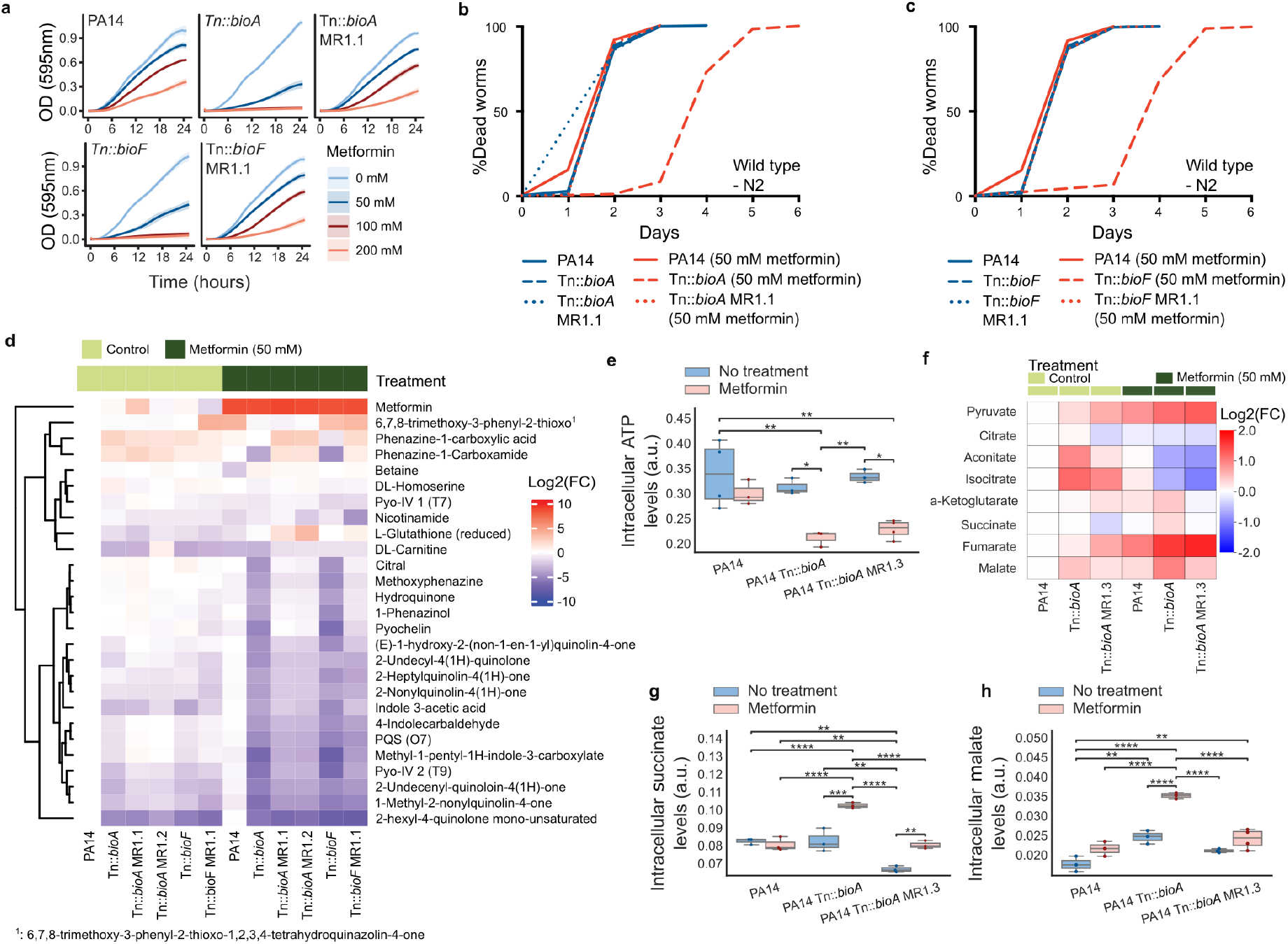
MR mutation induces changes within virulence and metabolism. **a,** Growth curves for PA14, PA14 Tn::*bioA*, PA14 Tn::*bioA* MR1.1, PA14 Tn::*bioF*, and PA14 Tn::*bioF* MR1.1 with increasing concentrations of metformin. (*n* = 3). **b, c,** Survival of *C. elegans* when exposed to PA14 Tn::*bioA* MR1.1 (*n* = 271-363) (**b**) or PA14 Tn::*bioF* MR1.1 (*n* = 336-396) (**c**) and treated with metformin. **d,** Log_2_(FC) data for untargeted metabolomics of PA14, Tn::*bioA*, Tn::*bioF* and their respective MR mutants with and without metformin vs. PA14 no treatment control (*n* = 3). **e, g, h,** Intracellular levels of ATP (**e**), Succinate **(g),** and Malate (**h**), for PA14, Tn::*bioA*, and Tn::*bioA* MR1.3. *n* represented by dot points. (ns: p > 0.05, *: 0.01 < p <= 0.05, **: 0.001< p <= 0.01, ***: 0.0001 < p <= 0.001, ****: p <= 0.0001) **f,** Log_2_(FC) values for TCA cycle intermediates for PA14, Tn::*bioA*, and Tn::*bioA* MR1.3, with and without metformin vs. PA14 no treatment control (*n* = 3-4). All statistics are provided in Extended Data Table 1 and 8.

**Extended Data Figure 6:**
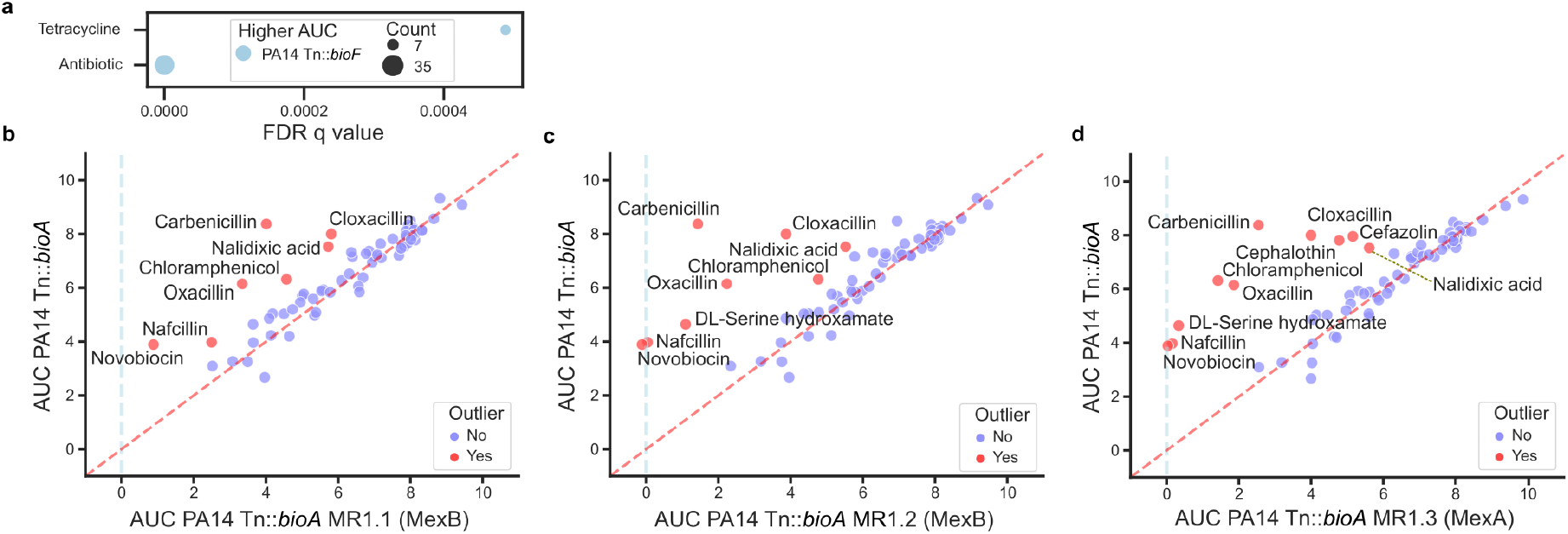
Metformin resistance leads to antibiotic sensitivity in biotin mutants of PA14. **a,** Enrichment of compounds with significant differences in AUC for the BIOLOG screen between PA14 Tn::*bioF* and PA14 Tn::*bioF* MR1.1. **b-d,** Subset BIOLOG chemical sensitivity screen comparing the average AUC for growth of PA14 Tn::*bioA* and PA14 Tn::*bioA* MR1.1 (**b**), PA14 Tn::*bioA* MR1.2 **(c**), or PA14 Tn::*bioA* MR1.3 (**d**), for all replicates within all concentrations of different chemical perturbations. (*n* = 1). All statistics are provided in Extended Data Table 9.

To evaluate the specificity of the MexAB-OprM mutation response to the combination of biotin synthesis disruption and metformin, we tested the ability of other mutants of PA14, that were identified in our high-throughput and survival screens, to gain metformin resistance (Fig. 1d,f and Extended Data Fig. 2a,b,c,e,f,j,l,p). Some strains, including PA14 Tn::*pyrE* developed suppressor colonies that exhibited improved growth in metformin compared to their parent strains (Extended Data Fig. 7a, Extended Data Table 10). However, enhanced growth was also observed in the absence of treatment. To determine whether this phenotype reflected true metformin resistance, we performed MIC assays. The results showed no significant differences in metformin resistance between suppressor colonies and their parent strains (Extended Data Fig. 7b, Extended Data Table 10). Despite the lack of metformin resistance, mutations abrogating the metformin-independent increase in survival of *C. elegans* exposed to PA14 Tn::*pyrE* did arise during the suppressor screen (Extended Data Fig. 7c, Extended Data Table 1). Therefore, MexAB-OprM mediated metformin resistance is highly specific to biotin deficient mutants of PA14.

Having confirmed specificity for biotin synthesis disruption, we investigated whether alternative clinically relevant biguanides would equally synergise with biotin limitation. MIC assays for PA14, PA14 biotin deficient mutants, and their MR suppressors were performed with phenformin and moroxydine. Unlike metformin, phenformin did not synergise with biotin synthesis disruption, with all strains tested demonstrating an MIC of 12.5 mM. However, moroxydine showed a conserved response to metformin treatment, with both PA14 Tn::*bioA* and PA14 Tn::*bioF* showing increased sensitivity to growth inhibition, which was not observed in metformin resistant strains, suggesting a conserved molecular mechanism (Extended Data Fig. 7d, Extended Data Table 10). Thus, the effect of moroxydine on survival was also examined and showed that, like for metformin, biotin inhibition and moroxydine extend survival (Extended Data Fig. 7e, Extended Data Table 1). Likewise, suppressor mutations also developed (Extended Data Fig. 7f,g, Extended Data Table 10). Similar to the MR mutants, *oprM* mutations were identified in moroxydine resistant (moxR) mutants. Therefore, given the similar MIC and survival results, biguanides are excellent molecular tools for non-antibiotic induction of AMR reversal.

Finally, we tested the specificity of this AMR reversal mechanism for other Gram-negative ESKAPE pathogens, namely biotin mutants of *E. coli* and *Acinetobacter baumannii*. *E. coli* and *A. baumannii* strains showed no sensitivity to metformin with the loss of biotin synthesis (Extended Data Fig. 7h,i, Extended Data Table 10), suggesting that biotin-mediated metformin resistance is likely PA specific among Gram-negative ESKAPE pathogens.

**Extended Data Figure 7:**
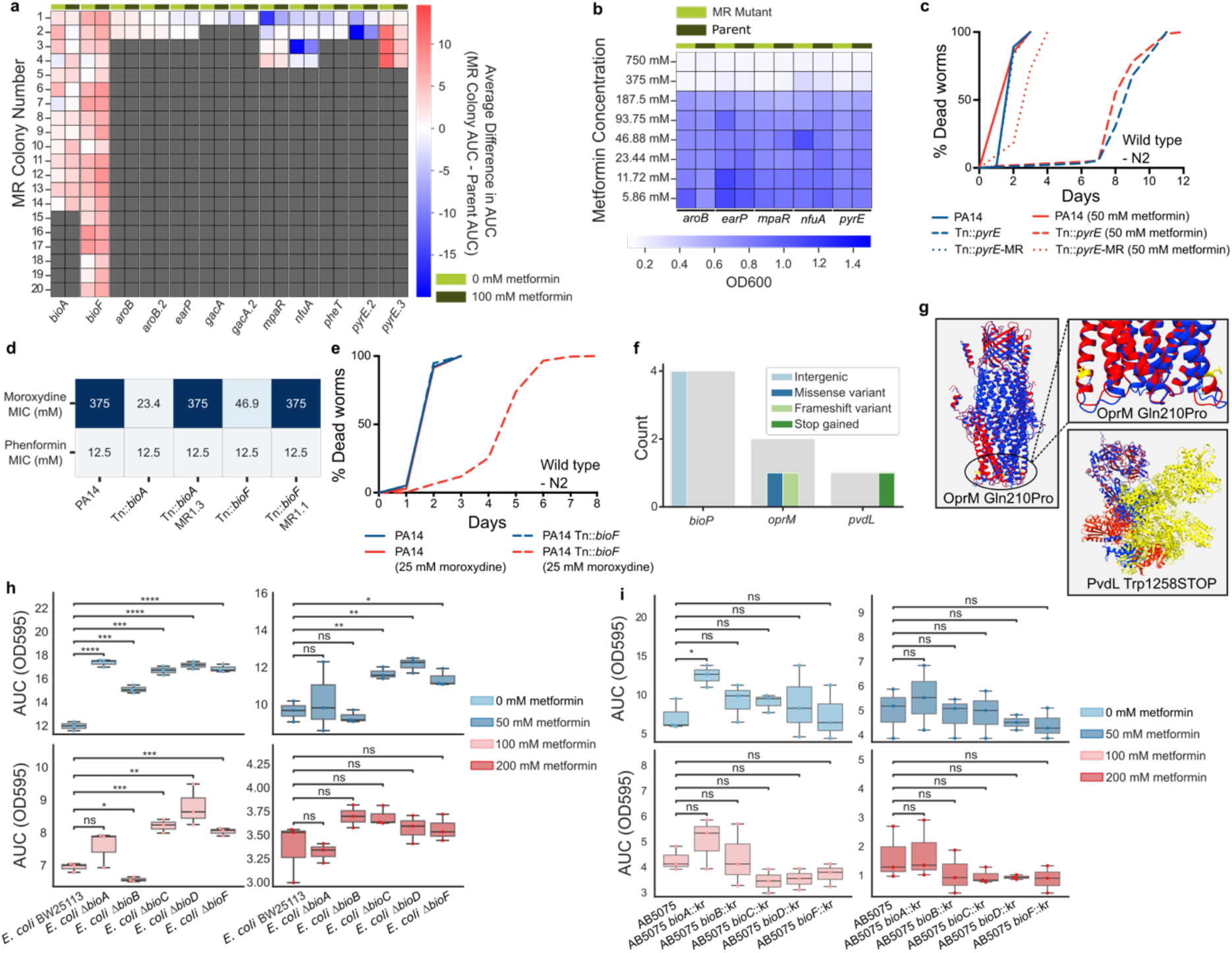
AMR reversing mutations are unique to biotin mutants, and some but not all alternative biguanides. **a,** Difference in AUC between potential MR colonies from the extended screen and their parents, for growth with and without metformin. (*n* = 1). **b,** MIC assay of potential MR colonies from the extended screen and their parents for metformin. (*n* = 1). **c,** Survival of *C. elegans* exposed to potential alternative MR mutant PA14 Tn::*pyrE-*MR with and without metformin treatment (*n* = 91-161). **d**, MIC of metformin resistant mutants for metformin analogues phenformin and moroxydine. (*n* = 3). **e,** Survival of *C. elegans* exposed to PA14 or PA14 Tn::*bioF* with and without moroxydine treatment (CPH p < 0.0001, *n* = 229-315). All survival statistics are provided in Extended Data Table 1. **f,** Location and type of moroxydine resistance (mox^R^) conferring mutations within PA14 Tn::*bioA* and PA14 Tn::*bioF.* **g,** AlphaFold structures of WT OprM and mox^R^ missense mutant OprM, and WT PvdL and mox^R^ premature stop mutant PvdL. WT structures (red) are overlayed with mox^R^ Structures (blue). Yellow segments represent the effected WT residues. **h,i,** AUC values for growth curves of *E. coli* BW25113 (**h**) or *A. baumannii* AB5075 (**i**) and their biotin synthesis mutants in increasing concentrations of metformin. (ns: p > 0.05, *: 0.01 < p <= 0.05, **: 0.001< p <= 0.01, ***: 0.0001 < p <= 0.001, ****: p <= 0.0001). (*n* = 3). All statistics are provided in Extended Data Table 10.

### Metformin and biotin synthesis disruption effects are conserved within clinical isolates

Metformin combination with biotin synthesis disruption displayed multiple beneficial attributes, decreasing virulence and microbial growth, as well as leading to multidrug sensitivity within the lab strain UCBPP-PA14. To test for conservation of these mechanisms among clinically relevant *P. aeruginosa* strains, we investigated 15 clinical isolates and disrupted biotin synthesis gene *bioA* within four, resulting in PA.CI-1 Δ*bioA*, PA.CI-3 Δ*bioA*, PA.CI-9 Δ*bioA*, and PA.CI-15 Δ*bioA*. These four were chosen for their increased ease of genetic manipulation and their differences in AMR profiles (Fig. 4a, Extended Data Table 12).

**Figure 4:**
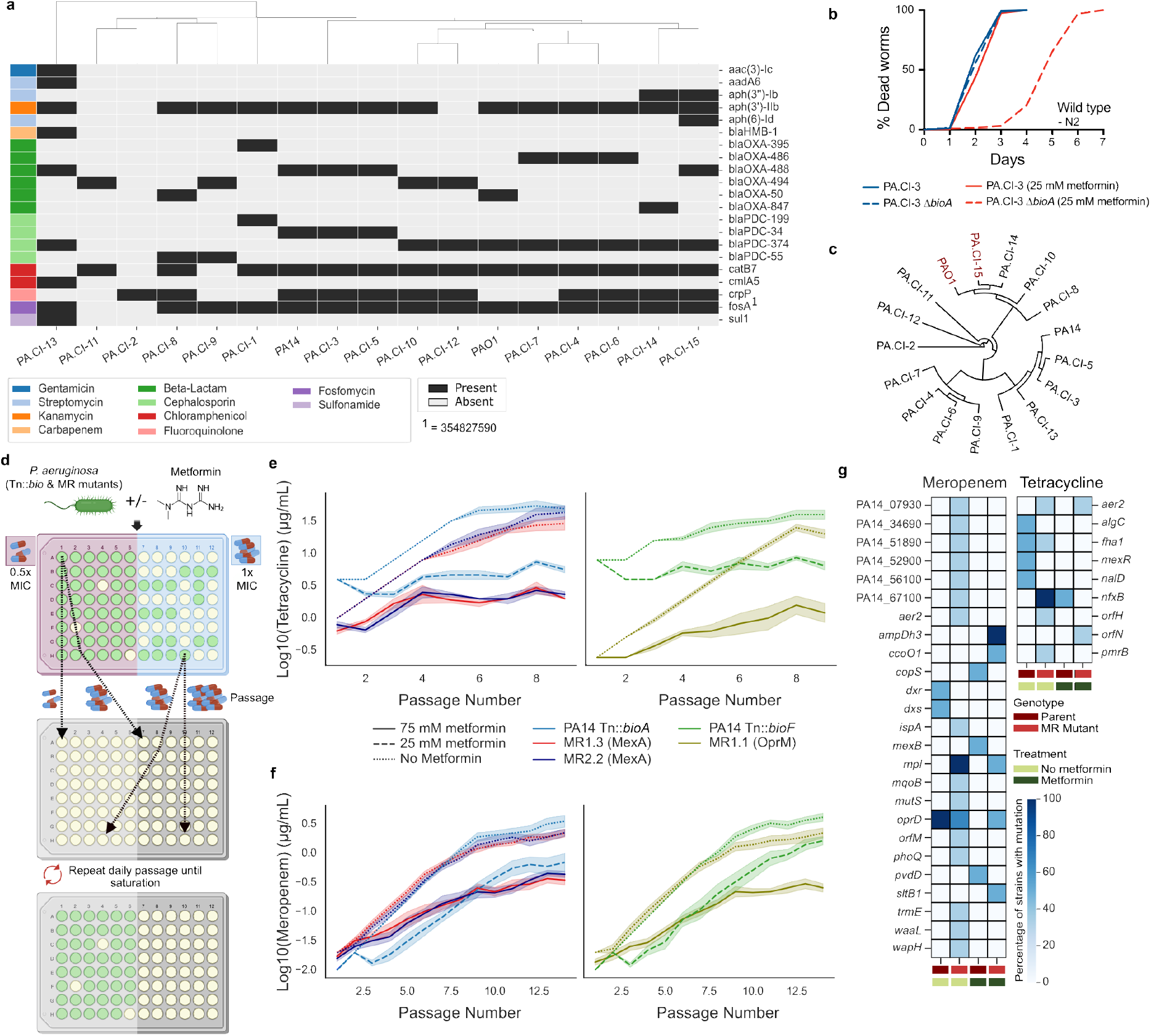
Metformin restricts the re-development of resistance to tetracycline and meropenem. **a,** Resistance genes present within clinical isolates sequenced within the study. **b,** *C. elegans* survival exposed to PA.CI-3 and PA.CI-3 Δ*bioA* with and without metformin (p < 0.0001, *n* = 265-361). All survival statistics are provided in Extended Data Table 1. **c,** Phylogenetic tree of clinical isolates. **d,** Schematic of the directed evolution experiment process. **e,f,** Evolution curve for resistance to tetracycline (**e**) or meropenem (**f**), representing the average highest concentration of drug which allowed growth per passage for PA14 Tn::*bioA* mutants (left) and PA14 Tn::*bioF* mutants (right) in the presence or absence of metformin. (*n* = 9). **g,** Percentage of strains with non-synonymous mutations in specific genes following the meropenem or tetracycline evolution experiments for end-point strains of PA14 Tn::*bioA*, PA14 Tn::*bioF*, PA14 Tn::*bioA* MR1.3, PA14 Tn::*bioA* MR2.2, and PA14 Tn::*bioF* MR1.1 with and without metformin. (Mer/Tet-Parent-No metformin: *n* = 2, Mer/Tet-Parent-Metformin: *n* = 2, Mer/Tet-MR Mutant-No metformin: *n* = 3, Mer-MR Mutant-Metformin: *n* = 2, Tet-MR Mutant-Metformin: *n* = 3). Intergenic mutations are not displayed. All statistics are provided in Extended Data Table 12.

First, we confirmed susceptibility to metformin with biotin disruption for each biotin-limited clinical isolate (BLCI). Concurrent with the PA14 data, a reduction in growth was displayed when exposed to metformin, regardless of the strain. Furthermore, exogenous biotin was sufficient to at least partially restore growth (Extended Data Fig. 8a, Extended Data Table 11). Interestingly, BLCIs were approximately 10-fold more susceptible to metformin than PA14 Tn::*bioA*, despite WT clinical isolates having similar MIC values to PA14 WT (Extended Data Fig. 8b, Extended Data Table 11). This is likely due to an increased biotin requirement within clinical isolates, since biotin supplementation aided the growth of PA.CI-1 Δ*bioA*, PA.CI-3 Δ*bioA*, and PA.CI-9 Δ*bioA* in the absence of metformin.

**Extended Data Figure 8:**
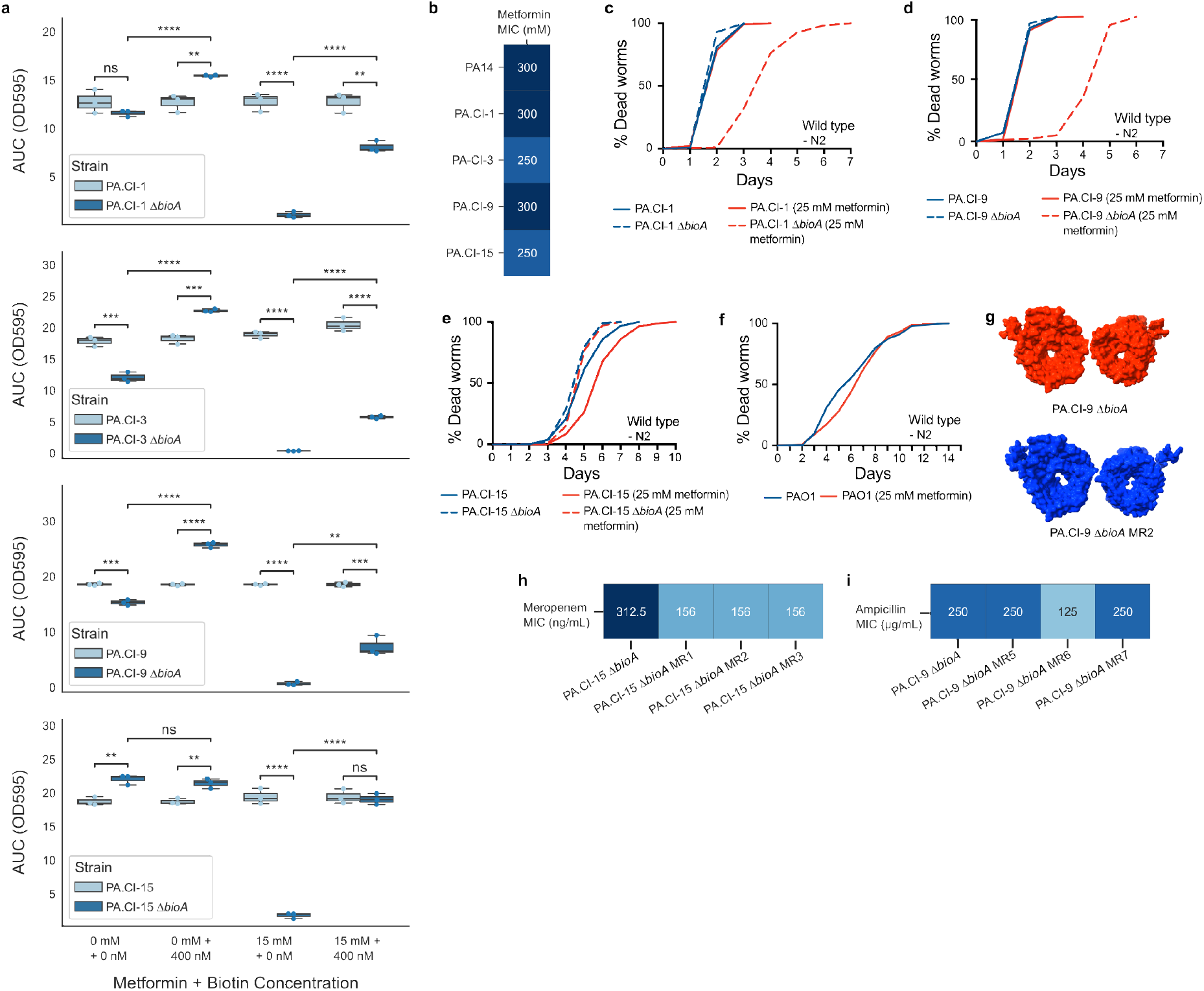
Metformin reduces growth and virulence of biotin limited *P. aeruginosa* clinical isolates. **a,** AUC values for growth of clinical isolates of PA and their biotin biosynthesis limited mutants (BLCIs) with and without metformin (15 mM) and/or biotin supplementation (400 nM). (ns: p > 0.05, *: 0.01 < p <= 0.05, **: 0.001< p <= 0.01, ***: 0.0001 < p <= 0.001, ****: p <= 0.0001), (*n* = 3). **b**, MICs of metformin against clinical isolates. (*n* = 3). **c-e,f,** *C. elegans* survival exposed to clinical isolates 1 (CPH p < 0.0001, *n* = 230-274), 9 (p < 0.001, *n* = 228-278) and 15, and BLCIs (CPH p = 0,0036, *n* = 258-290) (**c-e**), and PAO1 (**f**), and treatment with 25 mM metformin supplementation (*n* = 521-651). All survival statistics are provided in Extended Data Table 1. **g,** AlphaFold 3 structures of OprQ from WT PA.CI-9 Δ*bioA* (red) and metformin resistant mutant PA.CI-9 Δ*bioA* MR2 (blue). **h,i,** MIC of meropenem against PA.CI-15 Δ*bioA* (**h**) and ampicillin PA.CI-9 Δ*bioA* (**i**) and their MR mutants. (*n* = 3). All statistics are provided in Extended Data Table 11.

Next, we tested the *in vivo* efficacy of metformin against BLCIs via *C. elegans* survival assays. Complementary results were achieved for PA.CI-1 Δ*bioA*, PA.CI-3 Δ*bioA*, and PA.CI-9 Δ*bioA* with a significant extension of survival observed for N2 worms grown on these strains and treated with metformin. Survival on PA.CI-1 Δ*bioA* was similar to PA14 Tn::*bioA* with metformin treatment, yet at half the dose of metformin (Extended Data Fig. 8c, Extended Data Table 1) and a further extension was observed for PA.CI-3 Δ*bioA*, and PA.CI-9 Δ*bioA* in the presence of metformin (Fig. 4b, Extended Data Fig. 8d, Extended Data Table 1). Intriguingly, phylogenetically distant isolates PA.CI-15 WT and PAO1 to PA14 and the other clinical isolates, (Fig. 4c, Extended Data Table 12) are less virulent to *C. elegans* with or without metformin treatment. The loss of biotin, with or without metformin, worsened mean survival for PA.CI-15 with metformin (Extended Data Fig. 8e, Extended Data Table 1). With the BLCI data complementing the PA14 data, we subsequently produced metformin resistant BLCI (BLCI-MR) mutants. BLCI-MR mutants were obtained for all BLCIs and 9 were selected for sequencing for each strain. The suppressor mutations were mapped and 91.7% (33/36) BLCI-MR mutants showed mutations within the MexAB-OprM efflux pump (Fig. 3f, Extended Data Table 7). Three mutants lacking MexAB-OprM mutations were identified. In PA.CI-1 Δ*bioA* MR7, a SNP occurred in the intergenic region upstream of *bioP*. PA.CI-1 Δ*bioA* MR9 acquired an insertion that restored a conserved hypothetical gene homologous to PA1450, which was absent in the parental genome. PA.CI-9 Δ*bioA* MR2 carried a deletion in *oprQ*, resulting in the shortening of loop L7; a pore-constricting region characteristic of OprD family porins^27^. AlphaFold 3 structural comparisons of PA.CI-9 Δ*bioA* and PA.CI-9 Δ*bioA* MR2 OprQ revealed widening of the pore due to the L7 deletion, potentially enhancing substrate influx (Extended Data Fig. 8g). As overexpression of *oprQ* has been linked to increased antibiotic susceptibility, increased pore widening may likewise affect drug permeability^28^.

We next tested all BLCI-MR mutants for increased susceptibility to known substrates of the MexAB-OprM efflux pump (Fig. 3g, Extended Data Table 7). All BLCI-MR mutants showed increased resistance to metformin compared to the wildtype, confirming their MR status. Except for PA.CI-1 Δ*bioA* MR7, all other BLCI-MR mutants showed significantly reduced growth in the presence of carbenicillin, chloramphenicol and tetracycline. For meropenem, PA.CI-1 Δ*bioA,* PA.CI-3 Δ*bioA*, and PA.CI-9 Δ*bioA* MR mutants, excluding PA.CI-1 Δ*bioA* MR7 and PA.CI-9 Δ*bioA* MR2 displayed increased sensitivity. However, PA.CI-15 Δ*bioA* MR mutants did not show changes in sensitivity to meropenem, likely reflecting the lower baseline sensitivity of the parent strain. Further, fine-tuned MIC assays for a subset of PA.CI-15 Δ*bioA* MR mutants revealed a 2-fold increase in sensitivity to meropenem (Extended Data Fig. 8h, Extended Data Table 11). For ampicillin, nine MR mutants showed increased sensitivity to ampicillin, with PA.CI-9 Δ*bioA* MR mutants appearing more resistant. Therefore, we performed validation MICs for ampicillin for a subset of PA.CI-9 Δ*bioA* MR mutants. PA.CI-9 Δ*bioA* MR mutants MR5, and MR7 did not differ in MIC compared to PA.CI-9 Δ*bioA* and PA.CI-9 Δ*bioA* MR6 showed a 2-fold increase in sensitivity (250 µg/mL vs 125 µg/mL) (Extended Data Fig. 8i, Extended Data Table 11). Overall, these data show that development of metformin resistance in clinical isolates of PA leads to reversal of antimicrobial resistance against several antibiotics.

### Metformin and biotin-depletion-induced antibiotic collateral sensitivity is maintained by the presence of metformin

Acquisition of MexAB-OprM mutations was a highly conserved response by biotin synthesis deficient PA isolates to metformin. Thus, through experimental evolution, we assessed whether antibiotic sensitivity was stable over time. Through daily passage, PA14 Tn::*bioA*, PA14 Tn::*bioF* and a subset of their MR mutant strains were exposed to strongly selective concentrations of tetracycline and meropenem antibiotic, with or without metformin (Fig. 4d). The maximal antibiotic concentration permitting growth was plotted over passage number (Fig. 4e-f, Extended Data Table 12), and endpoint resistance (Extended Data Fig. 9a-b, Extended Data Table 13) as well as evolution rates (i.e. slope of regression lines) were compared across conditions (Extended Data Fig. 9c-d, Extended Data Table 13).

Strikingly, metformin significantly lowered resistance levels to both antibiotics for all strains. For tetracycline plus metformin, MR strains displayed significantly lower end point concentrations than their parental strains (Extended Data Fig. 9a, Extended Data Table 13). Comparable evolution rates suggest the lower endpoint could be due to the increased baseline tetracycline sensitivity of the MR strains, suggesting that metformin can limit tetracycline resistance independent of genotype (Extended Data Fig. 9c, Extended Data Table 13). Regardless, MR mutants retain their increased sensitivity with metformin exposure. Sequencing of end-point mutants revealed an alteration of tetracycline-related mutation acquisition with metformin exposure. The parents and MR mutant strains without metformin gained mutations within common efflux-based AMR determining genes, such as *mexR*, *nalD* and *nfxB* ^29^. However, mutations did not occur in these genes for MR mutants exposed to metformin, likely due to the detrimental effects of efflux pump expression with biotin deficiency and metformin treatment (Fig. 4g, Extended Data Table 12). PA14 Tn::*bioA* MR2.2 did however gain a frameshift mutation (Val50fs) in *orfN* with metformin exposure; a previously described early mutation in tigecycline resistance ^30,31^. However, unlike here, this *orfN* mutation is typically adjoined or followed by an *nfxB* mutation ^30^. This suggests that metformin limits the standard tetracycline evolution trajectory, which would explain the reduced end point resistance of MR mutants treated with metformin.

For meropenem, PA14 Tn::*bio*F MR1.1 evolved resistance more slowly and reached a significantly lower endpoint concentration than its parent under metformin treatment (Extended Data Fig. 9b,d, Extended Data Table 13). Notably, the metformin-sensitive parents required lower initial concentrations of meropenem, despite having a higher independent meropenem resistance. This was due to a synergy between metformin and meropenem, which drastically decreased the meropenem MIC for the metformin-sensitive parents when combined with metformin. Together, these findings show that OprM disruption synergises with metformin treatment for the slowing of meropenem resistance evolution while maintaining antibiotic hypersensitivity. Sequencing of endpoint meropenem strains also revealed suppression of mutation acquisition with metformin co-exposure. Mutations in 17 individual genes were developed for MR mutants without metformin exposure compared to 5 for metformin treated MR mutants. Despite this, mutations within AMR determining genes ^32–35^ *mpl* (Parent = 3/3, MR=1/2) and *oprD* (Parent = 2/3, MR=1/2) were achieved for both conditions, however, they were more prevalent without metformin exposure (Fig. 4g, Extended Data Table 12). Two AMR-inducing mutations were unique to MR mutants with metformin, *sltB1* (1/2) and *ampDh3* (2/2) (Fig. 4g, Extended Data Table 12). Both mutations are associated with an increase in the production of the beta-lactamase AmpC ^32–36^. Despite gaining these resistance mutations, the MR mutants exposed to metformin still had lower endpoint resistance. This suggests that the further mutations achieved without metformin exposure could be responsible for the increased resistance, and that metformin limits their acquisition.

**Extended Data Figure 9:**
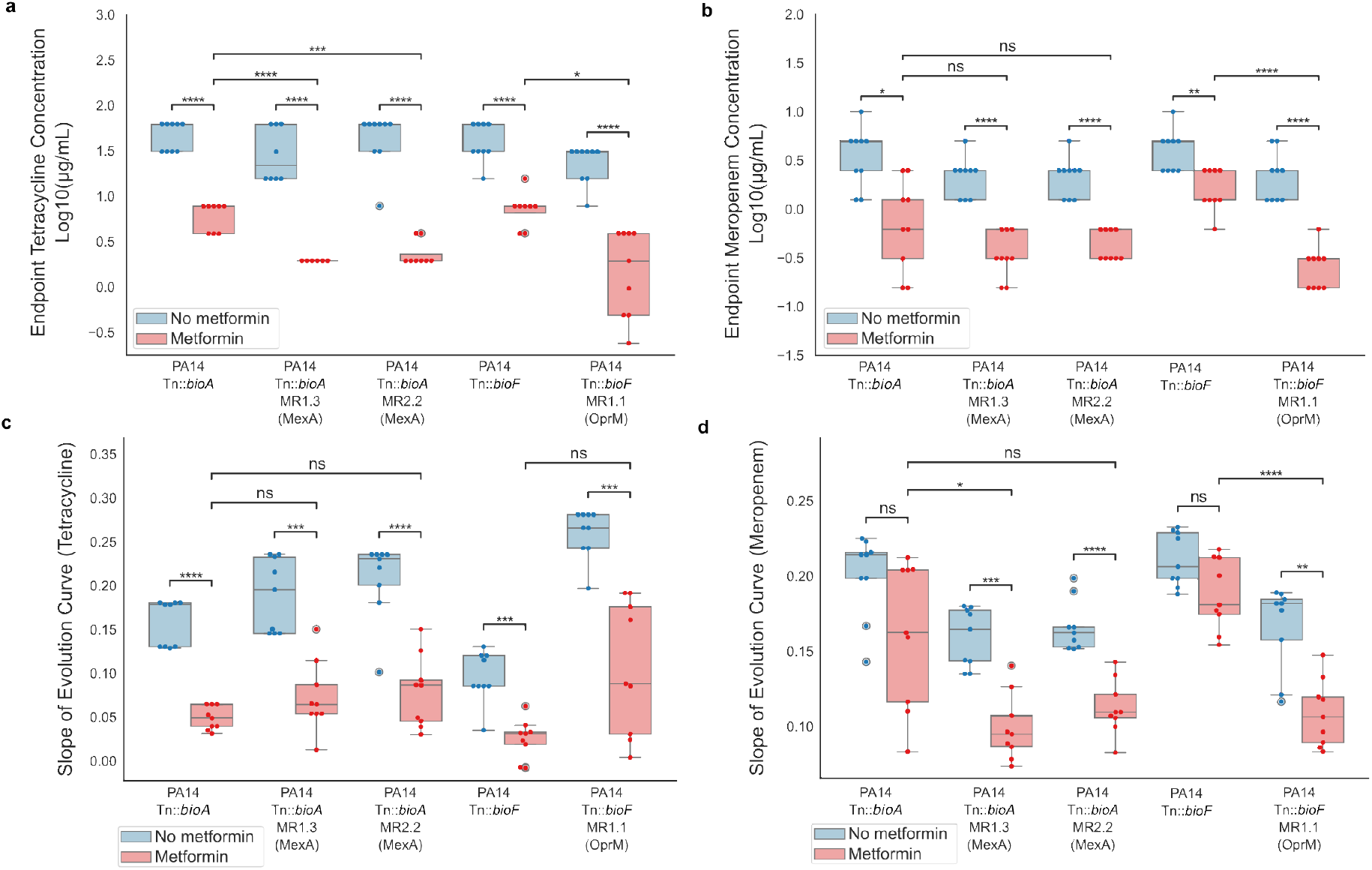
Metformin restricts the re-development of resistance to tetracycline and meropenem. **a,b,** Endpoint drug concentrations for each replicate of each strain, representing the highest concentration of tetracycline (a) or meropenem (**b**) that could be tolerated before plateau of resistance, with and without metformin. **c,d,** Slopes of the evolution curves for each replicate of each strain, indicating the rate of resistance evolution for tetracycline (**c**) or meropenem (**d**) with and without metformin. (*n* = 9), (ns: p > 0.05, *: 0.01 < p <= 0.05, **: 0.001< p <= 0.01, ***: 0.0001 < p <= 0.001, ****: p <= 0.0001). All statistics are provided in Extended Data Table 13.

## CONCLUSION

The reversal of antimicrobial resistance constitutes a critical frontier in global health and a pivotal objective in contemporary infectious disease research, aiming to restore the clinical efficacy of existing antimicrobial agents. As novel antibiotic discovery remains insufficient to offset escalating resistance rates, strategies enabling reversal and mitigation of AMR represent vital complements to drug innovation and stewardship efforts. Current reversal mechanisms involve interventions that disrupt or suppress established resistance determinants. These emerging approaches include the use of adjuvant compounds that inhibit resistance enzymes ^37^, bacteriophage therapy targeting resistant pathogens ^38^, CRISPR-based genetic tools^39^, and cycling or combinations of antibiotics, anti-infectives and natural products to induce collateral sensitivity (CS) ^40–48^. However, only one study has examined the role of non-antibiotic FDA-approved drugs for inducing CS. Recent work in *E. coli* showed that the development of resistance to non-antibiotics (i.e. sertraline), particularly through increased expression of drug efflux systems, can lead to cross-resistance to antibiotics ^49^, which may further aggravate AMR. Here, we present a unique conceptual two-pronged approach that targets virulence and forces reversal of AMR through collateral sensitivity. Namely, through the combination of biguanides with biotin synthesis inhibition. Collateral sensitivity was observed for 40/43 of metformin and 42/50 for biguanide-resistant mutants produced within the study due to loss-of-function mutations within the major efflux pump MexAB-OprM. Given the energy demands of MexAB-OprM for substrate excretion ^46^, these disruptive mutations may reduce pressure on the electron transport chain, reallocating energy resources for cell survival. Consistent with this, a study which performed Tn-seq on PA14 within energy-limiting conditions, found that insertions within the gene for the repressor of MexAB-OprM, *mexR*, induced a fitness disadvantage ^50^. Critically, the MexAB-OprM disrupted strains displayed increased sensitivity to multiple clinically relevant antibiotics, including the antibiotic of last resort meropenem. Furthermore, the rate of resistance re-development was heavily suppressed by biguanide presence, particularly for tetracycline. Common criticisms of CS studies focus on the effect of divergent genetic backgrounds upon the evolutionary trajectories of CS and highlight that repeatability both within and between strains must not be ignored ^48,51^. Importantly, CS within this study was conserved across multiple strains of PA, including clinical isolates with varying base AMR. Furthermore, CS was consistent across multiple replicates for each strain, suggesting a robust CS mechanism. Overall, the significance of reversing AMR through non-antibiotic approaches goes beyond direct therapeutic outcomes and, contrary to what was recently postulated, can preserve the effectiveness of existing antimicrobials ^49^.

Conventionally, development of antimicrobial drugs targeting other processes, such as central metabolism, has been frequently overlooked ^52,53^. Since central metabolism is highly interconnected and regulated, bacteria must consistently respond to their environment and balance the appropriate biosynthetic processes with energy generation for growth, survival and virulence ^54^. This is particularly true within infection environments where nutrients are typically limited, and a pathogen’s ability to utilise the available nutrient sources is paramount^54^. Since pathway essentiality can differ based on nutrient availability, understanding synthetic lethal gene-condition pairs could lead to the development of new metabolism-targeting antimicrobials. Here, we show that dual targeting of electron transport chain and biotin dependent metabolism creates a strong metabolic pressure on PA not achieved by each intervention alone. Consistent with the role of biguanides targeting the electron transport chain ^13,55–57^ and the role of biotin as a cofactor for different biotin-dependent carboxylases ^58^ involved in key energy processes such as fatty acid biosynthesis, the TCA cycle, gluconeogenesis, and amino acid metabolism, we observe significant changes in ATP production and rewiring of energy metabolism both at the proteome and metabolome levels. In particular, a reduction in TCA intermediates citrate, cis-aconitate and isocitrate could be explained by a blockage within oxaloacetate and acetyl-CoA production regulated by the biotin-dependent pyruvate carboxylase anapleurotic system. Consistent with an increased demand for alternative acetyl-CoA production, we observed a significant up-regulation of biotin-dependant carboxylases within the acetyl-CoA producing leucine (MCC) and acyclic terpene degradation (GCC) pathways with biotin limitation and metformin addition (Extended Data Fig. 4l, Extended Data Table 6). Our described mechanism for PA is consistent with observations for the antibacterial activity of non-antibiotics in *E. coli,* which is orthogonal to classic antibiotics ^49^. Critically, central metabolism has been extensively linked to pathogen virulence, as specific nutrients can act as signals for the expression of virulence factors ^59^. Here, we observe an important reduction in the arsenal of pathogenic mechanisms within PA, likely as an energy saving strategy under metabolic starvation for essential energy metabolites. This starvation dramatically reduced the pathogenic load and virulent nature of lab and clinical isolates of PA. Despite the conservation of metabolic pathways and enzymes across humans and bacteria, here we show that targeting of prokaryotic-unique metabolic pathways may be a sensible strategy. Significantly, the biotin synthesis pathway is absent in humans, making it an ideal target for drug discovery, due to the reduced likelihood of off-target effects.

Biotin plays pivotal roles in central metabolism in bacteria, and its biosynthesis is essential during infection with *A. baumannii*, *K. pneumoniae* and PA ^60,61^. Biotin biosynthesis inhibitors have been previously elucidated for the ESKAPE pathogens *E*. *coli* and *Mycobacterium tuberculosis*, such as MAC13772 (targeting both), acidomycin (*M. tb*), and amiclenomycin (*M. tb*). However, existing biotin biosynthesis inhibitors lack activity against PA ^61–63^, and no chemical inhibitors targeting its biotin pathway have yet been identified. The evolutionarily divergent biotin biosynthetic machinery in PA likely underpins species-specific advantages in metabolism, microbial competition, and virulence, while also creating a unique therapeutic vulnerability. Regardless, the discovery of inhibitors for other bacterial species, which effectively suppress biotin production and concomitantly reduce pathogenicity ^61–63^, suggests effective pathway druggability. The highly specific synthetic interaction between biguanides and biotin inhibition thus offers a promising strategy for precisely targeting PA virulence and reversing AMR. This approach provides a robust experimental and conceptual framework for developing similar virulence-targeting and AMR-reversal strategies against other ESKAPE pathogens minimizing the risk of disrupting commensal microbial communities or host physiology.

## METHODS

### Bacterial Strains Cultivation and Storage

All bacterial strains were stored within Lysogeny Broth (LB, Fisher Scientific) media supplemented with 15% glycerol (VWR) at -80°C. Overnight cultures were grown within LB supplemented with appropriate antibiotics at 37°C with aeration unless stated otherwise. Strains used are described within Extended Data Table 14.

### *C. elegans* strains and husbandry

Strains N2 wild type (CGC) and FGC86 sqIn82 [*argk-1p*::GFP + rol-6(su1006)]; wbmIs67 [*eft-3p*:: 3XFLAG::wrmScarlet::*unc-54* 3’UTR*wbmIs65] were used. Husbandry of *C. elegans* worms was performed on nematode growth media seeded with 120 µl of an overnight culture *E. coli* OP50 grown for a minimum of 2 days at 20°C, as previously established ^64^. Strains used are described within Extended Data Table 14.

### Nematode experimental conditions

The different media prepared for nematode cultivation included standard NGM, slow-killing (SK) media, and LB-NGM media. Standard NGM media contained: sodium chloride (3 gr/L, Fisher Scientific), Gibco™ Bacto™ peptone (2.5 gr/L, Fisher Scientific), BD Bacto™ agar (17 gr/L, BD), MgSO_4_ (1 mL/L Sigma-Aldrich), CaCl_2_ (1 mL/L, BioXtra), KH_2_PO_4_ (25 mL/L, Sigma-Aldrich) and cholesterol (1 mL/L, Sigma-Aldrich). In SK media, Gibco™ Bacto™ peptone quantity was increased to 4 gr/L, and in LB-NGM, bactopeptone was substituted for 4 gr/L of granulated LB. Where indicated, metformin (12.5, 25, 50, 100 mM, LKT Laboratories), or moroxydine (25 mM, TCI Chemicals) was supplemented to molten agar, or an equivalent volume of sterile dH_2_O. For the high-throughput screen, 96-square multi-well plates (Whatman) were prepared with LB-NGM media supplemented with 50 mM of metformin.

For all experiments worms were only used if consistently kept at optimal condition of 20°C for three generations. Worms were synchronised following standard alkaline hypochlorite (Thermo Scientific) treatment of gravid adult worms to obtain eggs. Eggs were allowed to hatch into L1 larvae in M9 media for a minimum of 12 hours. Larvae were placed on NGM with *E. coli* OP50 to continue development until L4-young adult worms.

### Preparation of *P. aeruginosa* for nematode experiments

Experimental set up of plates for *C. elegans* assay with PA was adapted from previous methods ^17,65^. Apart from the survival assay assessing the effects of three different media compositions, survivals were strictly performed on LB-NGM. A total of 50 µl of overnight cultures of PA strains of interest were seeded on LB-NGM plates (6 cm, Corning Life Sciences) supplemented with and without metformin and spread with an L-shaped glass spreader to cover a majority of the agar surface. Plates were incubated for 24 h at 37°C, followed by another 24 h at 25°C. Water reservoirs were added to provide optimal humidity for microbial growth. Around 2 h prior to worm addition, 5-fluoro-2-deoxyuridine (FUdR, 100 µM, Thermo Scientific) was topically added to each plate to prevent egg hatching.

Overnight cultures of the PA14 Transposon Insertion Library were added to the 96-square multi-well plates with LB-NGM media and metformin, by pipetting 5 µl of overnight culture per well with the VIAFLO 96 (Integra). Due to the reduction of plate surface for PA strains to grow, plates were incubated for 24 h at 25°C, followed by 24 h at 20°C with water reservoirs.

### *C. elegans* survival assays

Synchronised young adult worms were transferred to survival conditions through collection with M9 media. If collected worms showed turbid M9 due to presence of OP50, the worms were washed until turbidity was nearly removed. Worms were concentrated to reach a total amount of 30-60 worms per survival plate. Worms were incubated at 25°C for the entire survival duration. Unless stated otherwise, worms were scored for dead or alive every 24 hours. Worms were transferred to fresh survival plates on day 2 if worms were still alive. All survivals performed were pooled from three independent biological replicates, each containing three technical replicates with 30 worms on average. Survival results were processed in GraphPad Prism (Version 10) and statistical analysis including log rank test and Cox proportional hazards (CPH) were performed in JMP 18 (SAS Institute).

### *C. elegans* transcriptomics

RNA sequencing was performed on *C. elegans* N2 worms from NGM or LB-NGM plates, prepared with described PA14 strains in 4 independent biological replicates. Around 300-400 synchronised L4 larvae were added per plate, and a total of 4 technical replicates were prepared per condition to be able to collect around 1000 worms per condition after 24 hr incubation. Worm collection and RNA extraction was performed as previously described with adaptations^9^. Worms were collected in sterile nuclease free water for transfer to 2 mL lysing matrix D tubes (MP Biomedicals) of which the lysing beads were removed prior to worm addition. Worms were washed three times in sterile nuclease free water (Thermo Scientific), by waiting until worms sunk to the bottom of the tubes, carefully removing the supernatant and adding new nuclease free water. After the final washing, as much supernatant was removed as possible without disturbing the pellet of worms. Tubes were placed on ice to shrink the pellet further for supernatant removal. The lysing beads were returned to the tubes with 350 µL of TRIzol (Zymo research). Tubes were flash frozen in liquid nitrogen and stored at -80°C.

To extract RNA, frozen samples were vortexed until thaw, followed by flash freezing in liquid nitrogen. This was repeated twice as observation under the microscope still showed intact worm bodies after the first time. For the final thawing, samples were vortexed for 10 minutes using intervals of 2 minutes of vortexing followed by 30 seconds chilling on ice. A fixed volume of 280 µL per sample was transferred to RNAse free tubes. An equal volume of 280 µL 100% ethanol was added per sample, followed by RNA extraction using the Direct-zol RNA Miniprep kit (Zymo Research). RNA concentration and quality was measured and determined using both a NanoDrop 2000 spectrophotometer (Thermo Fisher Scientific) and a 2100 BioAnalyzer (Agilent). All samples were further processed by the LMS Genomics facility. Sequencing was performed on a NextSeq 2000 generating a minimum of 15 million 100bp read pairs per sample. Trimmomatic (v.0.39) was used to trim Illumina adapters and remove low quality reads, using a threshold of Q30 with a moving window of 10. Salmon (v.1.5.2) was used to quantify read counts with the reference genome of *C. elegans* from EnsemblGenomes (release 104) using the *–validateMappings* and *–gcBias* options. After quantification, the files were integrated in R with the package tximport (v. 1.18.0). Genes were annotated using AnnotationHub (v.3.2.0), followed by batch correction and statistical analysis with DESeq2 (v.1.30.1). Additional correction for visualisations was performed using limma package (v. 3.46.0) in R when necessary. To perform enrichment analysis, the web API from wormcat (v.2), String-db (api v.11-5) and the EnrichmentBrowser (2.20.7) package was used in R.

### Bacterial untargeted metabolomics

Overnight cultures of *P. aeruginosa* strains of interest were re-inoculated to an OD of 0.02 in 3 mL liquid LB-NGM supplemented with and without metformin (50 mM) for overnight treatment aerated at 37°C. The following day 1 mL per condition was centrifuged in Biorupter® Plus microtubes (Diagenode) or Eppendorf microtubes depending on the sonicator used. Samples were washed three times in sterile water, before complete removal of supernatant and flash freezing bacterial pellets in liquid nitrogen. Samples were saved at –80°C until further use. Further steps were adapted from previously described ^66^.

Flash frozen cells of respective samples were homogenized in 500 µL of ice-cold methanol/acetonitrile/water 5:3:2 (v/v/v) using the Precellys 24 homogenizer (Peqlab) at 6,400 rpm twice for 10 seconds with a 5-second pause. Per sample, 400 µL homogenate was extracted using a pre-cooled ThermoMixer (Eppendorf) at 4°C and 900 rpm for 20 minutes. After centrifugation (16,100 RCF, 20 min, 4 °C), the supernatant was dried using nitrogen gas, resuspended in 120 µl of water/acetonitrile 99:1 (v/v) and centrifuged again. After, 100 µL of the supernatant was transferred to autoinjector vials and subsequently analysed by Liquid Chromatography coupled to Electrospray Ionization High-Resolution Mass Spectrometry (LC-ESI-HRMS).

LC-ESI-HRMS analysis was performed by injecting 10 µl per sample onto a Core-Shell Kinetex C18 column (150 mm × 2.1 mm ID, 2.6 µm particle size, 100 Å pore size, Phenomenex) at 40°C and with detection using a Q Exactive HF quadrupole-orbitrap mass spectrometer (Thermo Scientific). The LC (Vanquish Horizon Binary UHPLC, Thermo Scientific) was operated at a flow rate of 0.3 ml/min with a mobile phase of water, 0.1% formic acid (solvent A) and acetonitrile, 0.1% formic acid (solvent B). Metabolites were eluted with the following gradient: initial, 1% B; 2 min, 1% B; 20 min, 100% B; 25 min, 100% B; 30 min, 1% B; 35 min, 1% B.

The mass spectrometer was equipped with a heated electrospray (HESI II) source. Samples were measured in random order by applying full scans while switching between the positive and negative ionization modes. The HESI source was operated with flow rates for sheath gas, auxiliary gas, and sweep gas of 55, 15 and 2, respectively. The spray voltage setting was 3.5/-3.5 kV, the capillary temperature 360°C, the S-lens RF level 50, and the auxiliary gas heater temperature 380°C. Full MS scans were acquired in the profile mode in the range of m/z 70-1000 with a resolution of 60,000, an Automatic Gain Control (AGC) target value of 3x10^6^ and a maximum injection time (IT) of 200 ms. Data-dependent MS/MS (ddMS^2^) experiments were performed on quality control samples (= pooled samples) - separately in positive and negative ionization mode - at the start and the end of each sequence. A “Top 5” method with an AGC target of 1×10^5^, a maximum IT of 50 ms, and a minimum AGC target of 1×10^3^ was used. The ions were isolated with a window of m/z 1.2, fragmented with stepped HCD energy NCE 15, 30, 45 and detected with a resolution of 30,000.

HRMS data was analysed with the metabolomics software Compound Discoverer® (Thermo Scientific, version 3.3.3.200) to perform searches based in the MS1 and MS2 levels of the mass spectrometry data against different databases. For the current project, two strategies were pursued; 1) general metabolomics and 2) specific secondary metabolites search, in both cases using Compound Discoverer®, but varying the databases to use. For general metabolomics an untargeted metabolomics workflow was used, which included different online and local databases search, such as: mzCloud, ChemSpider, Human metabolome database (HMDB), an in-house mzVault DB based on the Sigma Mass Spectrometry Metabolite Library (MSMLS), natural products atlas 2023 and Metabolika pathways database. Specific parameters for the identification were used, where the most important ones had the following values; retention time shift=0.5 min, mass tolerance=5ppm, preferred ions=[M+H]^+^ and [M-H]^-^. For specific secondary metabolites search, the same parameters above were used, but the search of potential targets was done by the use of a single database, which was locally built and included 53 known secondary metabolites produced by *Pseudomonas aeruginosa*, such as phenazines, quinolones, autoinducers, unsaturated carboxylic acids, terpenes, among others. Both analyses were considered and ultimately, data was filtered for annotated compounds, followed by selecting metabolites of interest for data visualisation. Data was transformed to log2 values and normalised for results of PA14 without metformin, in R to visualise with the ComplexHeatmap package.

### Targeted bacterial anionic metabolomics

LB-NGM broth with or without 50 mM metformin supplementation was inoculated 1:100 with overnight culture of PA14, PA14 Tn::*bioA* or PA14 Tn::*bioA* MR1.3. Cultures were grown to an OD600 ∼0.5 before centrifugation at 4000 rpm for 5 minutes. Supernatants were removed and pellets resuspended with the remainder before splitting in half. Cultures were centrifuged again and supernatant removed before flash freezing in liquid nitrogen. Levels of nucleotides, organic acids and sugar phosphates in the pellets were determined by Anion-Exchange Chromatography coupled to Electrospray Ionization High-Resolution Mass Spectrometry (IC-ESI-HRMS) using previously described procedures ^67,68^ with some modifications: *P. aeruginosa* cells were suspended in ice-cold methanol/acetonitrile/water 5:3:2 (v/v/v) using the Precellys 24 homogenizer (Peqlab) at 6,400 rpm twice for 10 seconds with a 5-sec pause. To 300 µl of homogenate, 25 µl of a mixture of isotope-labelled internal standards in Milli-Q water (50 µM 13C10-ATP, Sigma-Aldrich; 50 µM 13C6-D-glucose-6-phosphate (13C6-G6P) and 50 µM D4-succinic acid (D4-SUC), both Eurisotop) were added. The samples were extracted using a pre-cooled ThermoMixer (Eppendorf) at 4 °C and 900 rpm for 20 min. After centrifugation (16,100 RCF, 20 min, 4 °C), the supernatants were dried under a stream of nitrogen and resuspended in 100 µl of Milli-Q water. 80 µl of the supernatants were transferred to autoinjector vials and immediately analysed. IC-HRMS analysis was performed using a Dionex Integrion RFIC system (Thermo Scientific) coupled to a Q Exactive HF quadrupole-orbitrap mass spectrometer (Thermo Scientific) as previously described ^67,68^. The exact m/z traces of the internal standards and endogenous metabolites were extracted and integrated using the TraceFinder 5.1 software (Thermo Scientific). Endogenous metabolites were quantified by normalizing their peak areas to those of the internal standards: 13C10-ATP was used for nucleotides, 13C6-G6P for sugar phosphates and D4-SUC for organic acids. The peak area ratios were normalized to the protein content determined by a protein assay with bicinchoninic acid in a duplicate of each sample. ‘Area ratio/µg protein’ values were subjected to z-score analysis to determine outliers amongst replicates for each strain and metabolite. Values with an absolute z-score >1.5, representing 1.5 standard deviations from the mean of all replicates for the given metabolite and strain, were removed as outliers. A two-way ANOVA and Tukey-HSD adhoc test was employed to determine significance differences between all strain-treatment pairs for each metabolite. Log_2_(FC) values were also calculated for all strain-condition pairs against the PA14 no metformin control and displayed as a heatmap.

### Bacterial proteomics

Similar to untargeted metabolomics, overnight cultures of *P. aeruginosa* strains of interest were re-inoculated to an OD of 0.02 in 4 mL liquid LB-NGM overnight growth aerated at 37°C. 2 mL bacteria were centrifuged, and pellets were flash frozen in liquid nitrogen for storage at - 80°C.

Proteins were extracted in 8M urea through sonication using Biorupter® Sonication at high amplitude with 30 second on and off cycles. Samples adjusted to 100 µg per sample after quantification using a standard Bradford assay. Protein samples were processed using an in-solution digestion procedure. Briefly, samples were sequentially reduced and alkylated at room temperature and in the dark, to final concentrations of 10 mM dithiothreitol (DTT) and 50 mM 2-chloroacetamide (2-CAM), respectively. Samples were diluted 8-fold with 20 mM HEPES (pH 8.0), reducing the urea concentration to 1 M. This was followed by addition of 2 µg of trypsin (Promega, V528A). Samples were incubated over-night at 37°C. The digestion was stopped by acidification with 10% trifluoroacetic acid (TFA) to a final concentration of 1% and protein digests were desalted using Glygen C18 spin tips (Glygen Corp, TT2C18.96). Tryptic peptides were eluted with 60% acetonitrile, 0.1% formic acid (FA). Eluents dried by vacuum centrifugation.

Dried tryptic digests were re-dissolved in 0.1% TFA and each sample injected at 2 µg on-column. Liquid chromatography-tandem mass spectrometry (LC-MS/MS) analysis was performed using an Ultimate 3000 RSLC nano liquid chromatography system (Thermo Scientific) coupled to a coupled to a Q-Exactive mass spectrometer (Thermo Scientific) via an EASY spray source (Thermo Scientific). For LC-MS/MS analysis re-dissolved peptides were injected and loaded onto a trap column (Acclaim PepMap 100 C18, 100μm × 2cm) for desalting and concentration at 8μL/min in 2% acetonitrile, 0.1% TFA. Peptides were then eluted on-line to an analytical column (Acclaim Pepmap RSLC C18, 75μm × 75cm) at a flow rate of 200 nL/min. Peptides were separated using a 90 minute gradient, 4-25% of buffer B for 60 minutes followed by 25-45% buffer B for another 30 minutes (composition of buffer B – 80% acetonitrile, 0.1% FA) and subsequent column conditioning and equilibration. Eluted peptides were analysed by the mass spectrometer operating in positive polarity using a data-dependent acquisition mode. Ions for fragmentation were determined from an initial MS1 survey scan at 70,000 resolution, followed by HCD fragmentation (Higher Energy Collision Induced Dissociation) of the top 15 most abundant ions and MS2 scans at 17,500 resolution. MS1 and MS2 scan AGC targets were set to 3e6 and 5e4 for maximum injection times of 50ms and 100ms respectively. A survey scan m/z range of 350 – 1800 was used, normalised collision energy set to 27%, charge exclusion enabled with unassigned and +1 charge states rejected and a minimal AGC target of 1e3. Dynamic exclusion was set to 45 seconds.

Data was processed using the MaxQuant software platform (v1.6.2.3), with database searches carried out by the in-built Andromeda search engine against the Uniprot *P. aeruginosa strain PA14* database (version 20200603, number of entries: 5,88). A reverse decoy search approach was used at a 1% false discovery rate (FDR) for both peptide spectrum matches and protein groups. Search parameters included: maximum missed cleavages set to 2, fixed modification of cysteine carbamidomethylation and variable modifications of methionine oxidation and protein N-terminal acetylation. Label-free quantification was enabled with an LFQ minimum ratio count of 1. ‘Match between runs’ function was used with match and alignment time limits of 0.7 and 20 minutes respectively. Data was transformed to normalised log2 values followed by student T-test comparisons. KEGG enrichment was performed in StringDB, visualised in R or python.

### Fluorescent imaging of transcriptional reporter *C. elegans*

For all fluorescent imaging, differential P*argk-1*::GFP and constitutive P*eft-3*::mCherry levels were measured. For the high-throughput fluorescent imaging of P*argk-1*::GFP worms on the PA14 Transposon Insertion Library, fluorescent levels were solely assessed with metformin (50 mM). Approximately 25-30 young adult worms were transferred per well via multi-dispensing in M9 liquid collection. In low-throughput conditions, approximately 25-30 young adult worms were manually transferred to plates supplemented with and without metformin containing PA strains of interest. All plates were incubated for 24 h incubation at 25°C. After 24 h, worms were anesthetised with 5% levamisole (Thermo Scientific). Imaging was performed on the Zeiss Axio Zoom V16 microscope system equipped with an AxioCam MRm camera operated by Zen 2 software (Zeiss) and GFP filterset (excitation: 450-490 nm; emission: 500-550 nm) or the RFP filterset (excitation: 559-585 nm; emission: 600-690 nm), as previously described^9^.

Low-throughput images were taken under a 63x objective and exported in CZI format. Fluorescence levels were quantified using ImageJ (Version 2.16.0). The mean fluorescence intensities were determined by calculating the intensity per pixel of the entire area containing groups of 8 worms, subtracted by background noise. Total fluorescent levels were quantified in Ilastik (Version 1.0) by training the pixel classification on recognizing groups of 8 worm bodies and selecting the total intensity in object classification ^69^. Statistical analysis was performed in GraphPad Prism using Two-way ANOVA.

High-throughput images were taken in an automated manner with multiple Z-stacks under a 25x objective. Images were exported in CZI format per 96-well plate containing images from both GFP and RFP channels. The most focussed image per well per plate was selected through a python pipeline applying methods Brenner’s and Gray-level variance ^70,71^. Images were further processed in Ilastik, by using the RFP images to train the software to recognize single worm bodies in pixel classification generating a mask per image with the pixel locations of the worms. The masks were then applied in the object classification to measure the mean fluorescent intensities of both GFP and RFP images per well per plate. Data was collated into one CSV file for analysis in R (R core team). As one biological replicate was performed in batches, all data was normalised for PA14 wild-type control. GFP fluorescence levels was further normalised for RFP fluorescence levels to account for minor difference in lamp exposure and area of measurement. PA strains for which we were unable to identify single worms, were excluded. 2 biological replicates were performed containing an average of 6 worms per condition, yet if there was only a single worm identified in one of the biological replicates, the condition was included. Enrichment of the top 2% of genes with the highest P*argk-1*::GFP fluorescence levels was performed in ShinyGo 0.82 using *Pseudomonas aeruginosa* (Taxonomy ID 287, source STRING version 11.5) as reference and a maximum of 200 pathways.

### Fluorescent imaging and intestinal bacterial measurements

All *P. aeruginosa* bacterial strains used were conjugated with pMF440 from the DAP auxotroph donor strain *E. coli* BW29427. The plasmid pMF440 (Addgene #62550) constitutively expresses mCherry and can be selected for with ampicillin (100 µg/mL). Due to *P. aeruginosa’s* natural resistance to ampicillin we doubled the ampicillin dose to 200 µg/mL where needed. Experimental set up for fluorescent imaging was similar to the method using transcriptional reporter worms but performed with *C. elegans* N2. Imaging and bacterial extraction from *C. elegans* intestine was adapted from previously described ^72^. Worms were topically sedated using 2% levamisole and collected off the plate with M9. Worms were washed 3 times in M9. After the final wash, worms were imaged in brightfield and the RFP filter on the Zeiss Axio Zoom V16 microscope. Images were overlayed in FIJI ImageJ (version 2.16.0) for visualisation. After imaging, worms were collected in 20 µL of M9 with 0.01% Triton X-100 (Sigma-Aldrich) to grind worms using a micro-pestle. After, 5 µL was transferred into a 96-well microplate containing 200 µL of LB with ampicillin (50 µg/mL, Fischer Scientific). Bacterial growth was measured at OD 595nm every 30 minutes for 20 h using the cytation3 plate reader and the Biotek Biospa 8.

### Bacterial growth assay

Bacterial strains were cultured overnight at 37°C. Cultures were diluted to a final OD600 of 0.001 in LB broth supplemented with either 0 mM, 15mM, 50 mM 100 mM or 200 mM metformin and either 0 nM or 400 nM biotin (Sigma-Aldrich). Optical density at 595nm was measured every 30 minutes for 24 h using the cytation3 plate reader and the Biotek Biospa 8, or the BMG Labtech CLARIOstar (*A. baumannii* strains). OD values were adjusted either by subtraction of the average of blank OD values (PA14 strains and *A. baumannii* strains) or by subtraction of the average starting OD values (clinical isolate and *E. coli* strains). Technical replicate OD values were averaged before comparison of biological replicates. AUC values of replicate curves were compared for significant differences using independent T-tests with Benjamini-Hochberg correction.

### Clinical isolate genome sequencing

Genomic DNA from the Clinical Isolates was extracted using the Qiagen DNeasy Ultraclean Microbial Kit, following the manufacturer’s instructions. Genomic DNA was sequenced via nanopore sequencing and resulting SAM files were converted to Fastq files using Picard package SamToFastq v2.9.0-1-gf5b9f50-SNAPSHOT. Fastq reads were filtered using Filtlong v0.2.1, removing the worst 8% of reads in terms of read bases and retaining only sequences > 1000 bp. Genomes were then assembled using Flye v2.9.4-b1799 with an asm coverage of 40, a genome size of 7,000,000 and three iterations. Genomes were then annotated using Prokka v1.14.6. The resulting fna files were investigated for AMR genes using ABRicate v1.0.1, and for production of a phylogenetic tree using MASHtree v1.4.6. Fna files for PAO1 (GCF_000006765.1) and UCBPP-PA14 (GCF_000014625.1) were downloaded from Pseudomonas.com, database version 22.1 ^73^.

### Generation of in-frame *bioA* and *bioB-H* operon deletion mutants

Gene knockout was performed as described in Klein et al. ^74^ using the pEXTK derivative of pEXG2 for the clinical isolates as described in Eggers et al. ^75^. Genomic DNA from the PA14 and Clinical Isolates PA.CI-1, PA.CI-3, PA.CI-9 and PA.CI-15 was extracted using the Qiagen DNeasy Ultraclean Microbial Kit, following the manufacturer’s instructions. For the clinical isolates, conjugation was alternatively performed with the DAP auxotrophic donor strain *E. coli* BW29427. BW29427 carrying the respective pEXTK::Δ*bioA* plasmid, were mixed with the relevant clinical isolate in a 2:1 ratio and incubated for 2.5 hours at 37°C. Ex-conjugants were selected for on 75 µg/mL gentamicin (Sigma-Aldrich) supplemented LB agar plates. Thereafter positive selection to obtain a second crossover was performed by incubating gentamicin resistant clones for 3 h in 5 ml LB medium containing IPTG (1 mM). Subsequently, bacteria were selected by streaking bacteria on LB agar plates containing 100 μg/ml azidothymidine (Acros Organics) and 1 mM IPTG. Bacteria were then tested for loss of gentamicin sensitivity and mutants were verified by PCR. Plasmids used are described within Extended Data Table 14.

### Generation of metformin resistant mutants

Overnight cultures of PA14 Tn::b*ioA*, PA14 Tn::*bioF*, PA.CI-1 Δ*bioA*, PA.CI-3 Δ*bioA*, PA.CI-9 Δ*bioA*, and PA.CI-15 Δ*bioA* were grown at 37°C before plating onto LB-NGM plate supplemented with 50 mM metformin (PA14) or 25 mM metformin (BLCIs). Plates were incubated overnight at 37°C before transferring to 25°C incubation until suppressor colonies formed. Suppressor colonies were collected in LB and cultured overnight for preparation of glycerol stocks. Metformin resistance was confirmed via growth assays, as described previously.

For the testing of alternative *argk*-1 inducing mutants of PA14, overnight cultures were prepared in 96 well plates and grown at 37°C. Square 96-well plates were prepared with LB/NGM agar and supplemented with either 50 mM, 100 mM , 150 mM or 200 mM metformin before inoculation with 10 µL of overnight culture. The overnight culture parent plate was supplemented with 12.5% glycerol and stored at -80°C. Cultures were grown overnight at 37°C before being transferred to 25°C and incubated for 4-6 days. Potential suppressor colonies were picked and transferred into 96-well LB broth plates and grown overnight at 37°C before supplementation with 12.5% glycerol and storage at -80°C. Resistance to metformin was first tested via growth assay in 0 mM or 100 mM metformin. Overnight culture of the 96 well plates for the suppressor mutants and parents were prepared from the glycerol stock plates and grown overnight at 37°C. Overnight cultures were diluted 1 in 10, and 1 µL of diluted culture was transferred from each well to corresponding wells within 96 well plates with either LB broth or LB broth + 100 mM metformin. Optical density at 595 nm was then measured every 30 minutes for 24 hours. Growth curves were smoothed using a Wiener filter before AUC values were calculated before calculating the difference between parent and mutant AUCs. Metformin resistance was further tested for a subset of suppressor colonies via broth microdilution MIC assay, as described below, with a concentration range of 750 mM to 5.85 mM. MIC OD value profiles were compared between parents and mutants using a student’s t-test and Bonferroni correction.

### Sequencing and mapping of resistance conferring mutations

Genomic DNA was extracted from each mutant and parent using the Qiagen DNeasy UltraClean Microbial Kit, following the manufacturer’s instructions. Clinical isolate MR mutants, as well as moroxydine resistant PA14 mutants, were sequenced and processed until assembled fasta files were achieved, as described above. For metformin resistant PA14 strains, genomic DNA was sequenced via Illumina sequencing. PA14 parent fastq paired end reads were assembled using Shovill v1.1.0. Parent genomes for both CIs and PA14 strains were then annotated using Prokka v1.14.6, using the compliant mode, to gain gbk files for use as reference sequences. Snippy v4.6.0 was then employed on the Flye assembled fasta files for the PA14 moxR mutants and the clinical isolate MR mutants, and on the paired end fastq files for the PA14 MR mutants to locate the resistance conferring mutations. The respective parent gbk files were used as the reference genomes. Protein sequences of MexA, MexB and OprM for the PA14 parents and PA14 MR mutants; the sequences for PA.CI-9 Δ*bioA* and PA.CI-9 Δ*bioA* MR2 OprQ; and the PA14 parent and moxR mutant OprM and PvdL sequences, were submitted to the AlphaFold 3 server for structure prediction. Mutated and parent proteins were overlayed using the Matchmaker tool within ChimeraX-1.8

### Broth microdilution minimum inhibitory concentration

PA14 Tn::*bioA*, PA14 Tn::*bioF*, PA.CI-9 Δ*bioA*, PA.CI-15 Δ*bioA* and their respective MR mutants were streaked on LB agar and grown overnight a 37°C. Day cultures of LB broth were inoculated with 5-6 colonies of respective bacterial strain and grown at 37°C for 5 h before dilution to an OD600 of 0.002. Carbenicillin (Sigma-Aldrich), chloramphenicol (Thermo Scientific), tetracycline (Sigma-Aldrich), and kanamycin (Fischer Scientific) were serially diluted within 96-well plates, from 500 µg/mL to 3.9 µg/mL, before addition of bacterial culture to a final OD of 0.001. Ampicillin was serially diluted from 1 mg/mL to 7.8 µg/mL, meropenem (TCI Chemicals) from 5 µg/mL to 0.039 µg/mL, metformin from 750 mM to 5.85 mM, moroxydine from 250 mM to 1.95 mM, and phenformin (Sigma-Aldrich) from 100 mM to 0.78 mM. Plates were grown at 37°C for 24 h before OD600 measurement. OD values were adjusted to the OD of the highest concentration of drug as a blank adjustment, since no growth occurred here for any strain in any drug. The OD values at each concentration of drug for three technical and three biological replicates for each strain was then averaged and growth was determined by an OD600 value > 0.1. The lowest drug concentration where OD600 < 0.1 was determined as the MIC.

### BIOLOG chemical sensitivity screening

Relevant bacterial strains were grown overnight at 37°C on LB agar plates. Several colonies were resuspended in IF-0a media, before OD adjustment to an OD600 of 0.07. Adjusted cultures were used to inoculate 0.01% Tetrazolium LB broth to a final OD600 of 0.0002, before addition to BIOLOG Microbial Phenotype plates PM11-20 and an additional plate named PMPenem, comprising of 88 no-drug controls and 4 concentrations of meropenem and biapenem. For PA14 Tn::*bioA*, PA14 Tn::*bioA* MR1.1, PA14 Tn::*bioA* MR1.2, and PA14 Tn::*bioA* MR1.3 only PM11, PM12, and PM20 were used. Optical density at 750 nm was then measured every 30 minutes for 24 h. OD750 values were blank adjusted using the mean OD value of their respective second to sixth time points, to account for colour differences introduced by the chemical stressors. Cross plate normalisation was performed by adjusting the OD values for each plate at each time point such that the OD values within the 40^th^ to 90^th^ percentiles of growth were equal to the average OD values across all plates at the given timepoint. Area under the curve (AUC) was calculated for each condition from the 3^rd^ to 24^th^ hour timepoints. The AUC scores were then averaged across the 4 concentrations of each compound within the BIOLOG library. T-tests were then performed for each condition between PA14 Tn::*bioF* and PA14 Tn::*bioF* MR1.1 and Cohen’s d was calculated to evaluate the effect size. Conditions with a *p*-value ≤ 0.05 and an absolute Cohen’s *d* > 1 were selected as significant and subjected to an enrichment analysis for molecule type. Enrichment was performed using LightGOEA and inputting a dictionary of molecule type terms for each condition. For PA14 Tn::*bioA*, PA14 Tn::*bioA* MR1.1, PA14 Tn::*bioA* MR1.2, and PA14 Tn::*bioA* MR1.3 conditions where the difference in AUC between the mutant and the parent was greater than an absolute Z-score threshold of 2 were considered outliers. This threshold represents AUC differences within the top and bottom 2.5% of the overall distribution as significant.

### BLCI MR mutant antibiotic sensitivity assay

Overnight cultures of BLCI parents and MR mutants were prepared in LB broth and grown overnight at 37°C. Cultures were diluted 1 in 10 in LB broth. 96 well plates were set up with 200 µL LB broth supplemented with either metformin (25 mM), meropenem (50 ng/mL), carbenicillin (5 µg/mL), tetracycline (2.5 g/mL), Chloramphenicol (15 µg/mL), or ampicillin (5 µg/mL). Plates were inoculated with 1 µL of appropriate diluted culture. Bacterial growth at 37°C was assessed via measurement of optical density at 595nm every 30 minutes for 24 h using the cytation3 plate reader and the Biotek Biospa 8. Three technical replicates were performed per strain per condition and three biological replicates undertaken. For each BLCI parent and its MR mutants AUC values (for each technical/biological replicate) were calculated, and a one-way ANOVA and pairwise Tukey-HSD analysis was performed on these values to determine statistically significant differences between parents and MR mutants. From this, the average difference in AUC between parents and mutants was calculated for each drug.

### Experimental evolution of antibiotic resistance

Overnight cultures of PA14 Tn::*bioA*, PA14 Tn::*bioF,* PA14 Tn::*bioA* MR1.3, PA14 Tn::*bioA* MR2.2, and PA14 Tn::*bioF* MR1.1 were prepared in LB medium and grown overnight at 37°C. 96 well plates were prepared with LB medium supplemented with either tetracycline or meropenem to a strain-dependent starting concentration, see Extended Data Table 12. Each 96 well plate constituted of two concentrations of drug, with a 2-fold difference in concentration. These plates were then supplemented with either LB or metformin to a final concentration of 25 mM for PA14 Tn::*bioA* and PA14 Tn::*bioF* and 75 mM for PA14 Tn::*bioA* MR1.3, PA14 Tn::*bioA* MR2.2, and PA14 Tn::*bioF* MR1.1. MR mutants received higher concentrations of metformin than parents to retain a selective pressure, since these had a higher metformin tolerance. Nine replicates for each strain were then added to both the lower and higher concentrations of antibiotic to a final OD600 of 0.001. Three wells were left un-inoculated and without antibiotic supplementation as contamination control wells. Plates were incubated at 37°C for 21 hours before OD600 was measured and the concentrations of antibiotics for the next passage were calculated. If growth was achieved in the highest concentration of antibiotic, then the concentration of antibiotic for this replicate was doubled for both concentrations within the new plate. If growth was only achieved in the lower concentration, concentrations were kept stable. Where growth was not achieved in either concentration, the concentration of antibiotic was halved and cultures from the last plate where the replicate grew were used for the passage. New 96 well plates with fresh media were prepared with the adjusted antibiotic concentrations and replicates were passaged (1 µL) to the new plate, taking from the well with the highest concentration of antibiotic with growth. This was repeated until increases in resistance stabilised. Glycerol was added to the final four plates for each condition and were stored at -80°C. OD600 values were adjusted to their respective plate blanks before being matched to their respective concentrations of drug. Where the OD600 value for the highest concentration was > 0.15 this concentration was taken for the log10 concentration of that passage number. If only the lower concentration had growth, then this concentration was taken. Where neither grew, the concentrations for the day before were taken from the highest concentration with growth. Averages of the log10 drug concentrations for each strain were then logged against the passage number. The slope of each replicate’s log_10_(concentration) vs passage day curve was calculated using the scipy.stats.linregress linear regression module. The statistical difference between slopes values, as well as endpoint concentrations, was calculated using a t-test and Bonferroni correction.

### Sequencing and mutation mapping of endpoint evolution experiment strains

Samples from one replicate of each strain for each condition were sub-cultured from the final passage plates on LB agar and grown overnight at 37°C. Genomic DNA was extracted from samples as previously described and sequenced via Nanopore sequencing. Bam files were converted to Fastq files and filtered as described above. The previously analysed fastq files for PA14 Tn::*bioA* MR1.3, PA14 Tn::*bioA* MR2.2, and PA14 Tn::*bioF* MR1.1 were annotated using Prokka v1.14.6 using the compliant mode and UCBPP-PA14 genome (GCF_000014625.1) as a reference, to gain gbk files for use as reference sequences, alongside those previously processed for PA14 Tn::*bioA* and PA14 Tn::*bioF*. The fastq files for the evolution samples were then mapped to their pre-evolution parents for mutations using Snippy v4.6.0. Counts for non-synonymous and non-intergenic mutations for each gene were calculated for each condition for parent strains or MR mutant strains. Percentage mutations were then calculated for each condition-strain set, by dividing the counts by the number of strains in the set and multiplying by 100. Where the presence of multiple mutations within the same gene for the same sample occurred only one mutation was counted.

## Supporting information

Extended Data Table content list

## ACKNOWLEDGEMENTS

FC was supported by the Wellcome Trust/Royal Society (102532/Z/12/Z and 102531/Z/13/A), the DFG German Research Foundation (EXC 2030 -390661388) and (IPFP-B02 Filipe Cabreiro) of the Center for Molecular Medicine Cologne. DMM was supported by the Leverhulme Trust (RPG-2022-299). We acknowledge computational resources and support provided by the Imperial College Research Computing Service (http://doi.org/10.14469/hpc/2232). HMD was supported by i-HEAD“ (PB-NRW-2022-ANTRAG-025) and the Bundesministerium für Forschung, Technologie und Raumfahrt (BMFTR) GO-Bio *initial* (03LWH0066). TF and TB were supported by German Research Foundation (DFG) Collaborative Research Centre (SFB) 1210. EB was supported by the DFG Cluster of Excellence EXC2124 ‘Controlling Microbes to Fight Infection’. MB was supported by a UKRI Future Leaders Fellowship [MR/V027204/1].

## AUTHOR CONTRIBUTIONS

HMD and JVDL have equally contributed, and order displayed is alphabetical. FC and MB conceptualized the research. AI and LG sequenced transcriptomes and genomes. IK, AM and HK performed proteomics identification and analysis. IS, JM and SB performed the metabolomic identification and analysis. DMM contributed to computational analysis. TB advised on evolution experiment design. EB contributed to the production of in-frame deletion mutants. TF, KS, HA, and LH contributed to experimental procedures. HMD, JVDL and FC wrote the manuscript with contributions from all authors. HMD, JVDL, MB and FC participated in the interpretation of the main findings. FC and MB supervised the research and secured funding. All authors read and approved the final manuscript.

