## Extended Data Table content list for "Restoring Antibiotic Sensitivity via Metabolic Pathway Manipulation in *Pseudomonas aeruginosa*"

**Extended Data Table Contents:**

extended-data-table1.xlsx – All survival data. Extended Data Figure 2 data.

extended-data-table2.xlsx – Figure 1 data

extended-data-table3.xlsx - Extended Data Figure 1 data

extended-data-table4.xlsx – Extended Data Figure 3 data

extended-data-table5.xlsx – Figure 2 data

extended-data-table6.xlsx – Extended Data Figure 4 data

extended-data-table7.xlsx – Figure 3 data

extended-data-table8.xlsx – Extended Data Figure 5 data

extended-data-table9.xlsx – Extended Data Figure S data

extended-data-table10.xlsx – Extended Data Figure 7 data

extended-data-table11.xlsx – Extended Data Figure 8 data

extended-data-table12.xlsx – Figure 4 data

extended-data-table13.xlsx – Extended Data Figure 9 data

extended-data-table14.xlsx – Strain and plasmids used within the study
